# Circadian control of lung inflammation in influenza infection

**DOI:** 10.1101/396556

**Authors:** Shaon Sengupta, Soon Yew Tang, Jill Devine, Soumyashant Nayak, Shirley Zhang, Alex Valenzuela, Carolina B. Lopez, Gregory Grant, Garret A. FitzGerald

## Abstract

Influenza is a leading cause of respiratory mortality and morbidity. While inflammation is necessary for fighting infection, a fine balance of anti-viral defense and host tolerance is necessary for recovery. Circadian rhythms have been known to modulate inflammation. However, the importance of diurnal variability in the timing of influenza infection is not well understood. Here we demonstrate that endogenous rhythms influence the cellular response to infection in bronchoalveolar lavage (BAL), the pulmonary transcriptomic profile and lesional histology. This time dependent variability does not reflect alterations in viral replication. Rather, we found that better time-dependent outcomes were associated with a preponderance of NK and NKT cells and lower proportion of monocytes in the lung. Thus, host tolerance, rather than viral burden underlies the diurnal gating of influenza induced lung injury.

**Significance statement:** Our work demonstrates the importance of circadian rhythms in influenza infection --a condition with significant public health implications. Our findings, which establish the role of the circadian rhythms in maintaining the balance between host tolerance pathways and anti-viral responses confers a new framework for evaluating the relevance of circadian influences on immunity.

## Introduction

Circadian rhythms constitute an innate anticipatory system with a 24-hr periodicity that helps the organism adapt to its surroundings and hence improve survival. At the molecular level, circadian rhythms are controlled by oscillating core clock genes, which regulate rhythmic expression of their downstream targets(1). Many physiological processes including immune responses(2-5) are under circadian regulation. Inflammation is a critical part of the immune response to influenza. An ineffective inflammatory response hinders viral clearance; however a high level of inflammation injures the host(6).(7) Due to its role in maintaining overall homeostasis, the circadian regulatory system may act to provide the host such a balance.

Although the role of circadian regulation in systemic viral infections(8) has been described using Murid herpes virus (MHV)(9) and Vesicular Stomatis Virus (VSV)(10), its purview for respiratory viruses is limited(11). Some *in vitro* work(9) and results of a cluster-randomized study testing the efficacy of influenza vaccine in older adults(12) have suggested a role for circadian rhythms in influenza A infection. Influenza infection is a leading cause of respiratory morbidity and mortality. However, the role of circadian rhythms in lung inflammation induced by influenza infection has yet to be systematically evaluated.

In this study, we tested whether the outcome of influenza A infection (IAV) was influenced by the time of infection, to understand whether the molecular clock might help balance host tolerance and the anti-viral response to IAV.

## Results

### Time of infection determines disease progression and survival

To ascertain whether the time of infection determines mortality and morbidity from influenza A infection, C57BL/6J mice (male and females in approximately equal numbers) were infected either just before the onset of their active phase/ lights off (active phase: ZT11) or just prior to the onset of their rest phase/lights-on (ZT23) with same dose of IAV (PR8 strain) intranasally (i.n.) (Fig. 1A). Animals were weighed and monitored daily for 2 weeks to record disease progression. Mice infected at ZT11 had significantly higher mortality (71% in ZT11 vs 15% at ZT23) than mice infected at ZT23 (Fig. 1B). Furthermore, after day 4 post-infection (p.i.) mice infected at ZT11 had more weight loss than mice infected at ZT23 (Fig 1C). These results were confirmed with mice on inverted light-dark cycle, where we were able to infect both groups simultaneously (data not shown). We also recorded clinical scores based on activity level, behavior and respiratory distress (Table. S1) and found higher scores consistent with increased morbidity and mortality (Fig. 1D) in mice infected at ZT11.

**Fig. 1:**
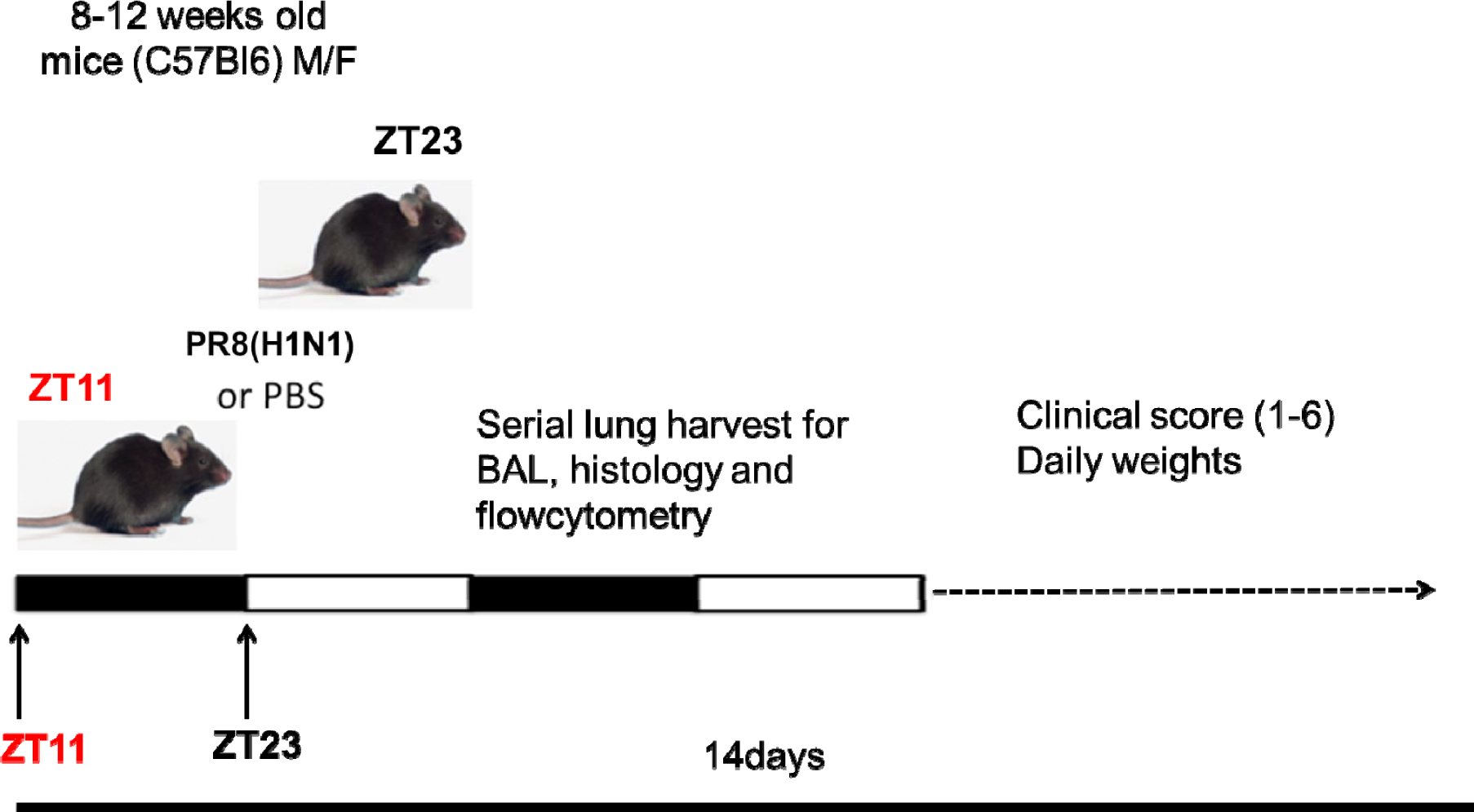
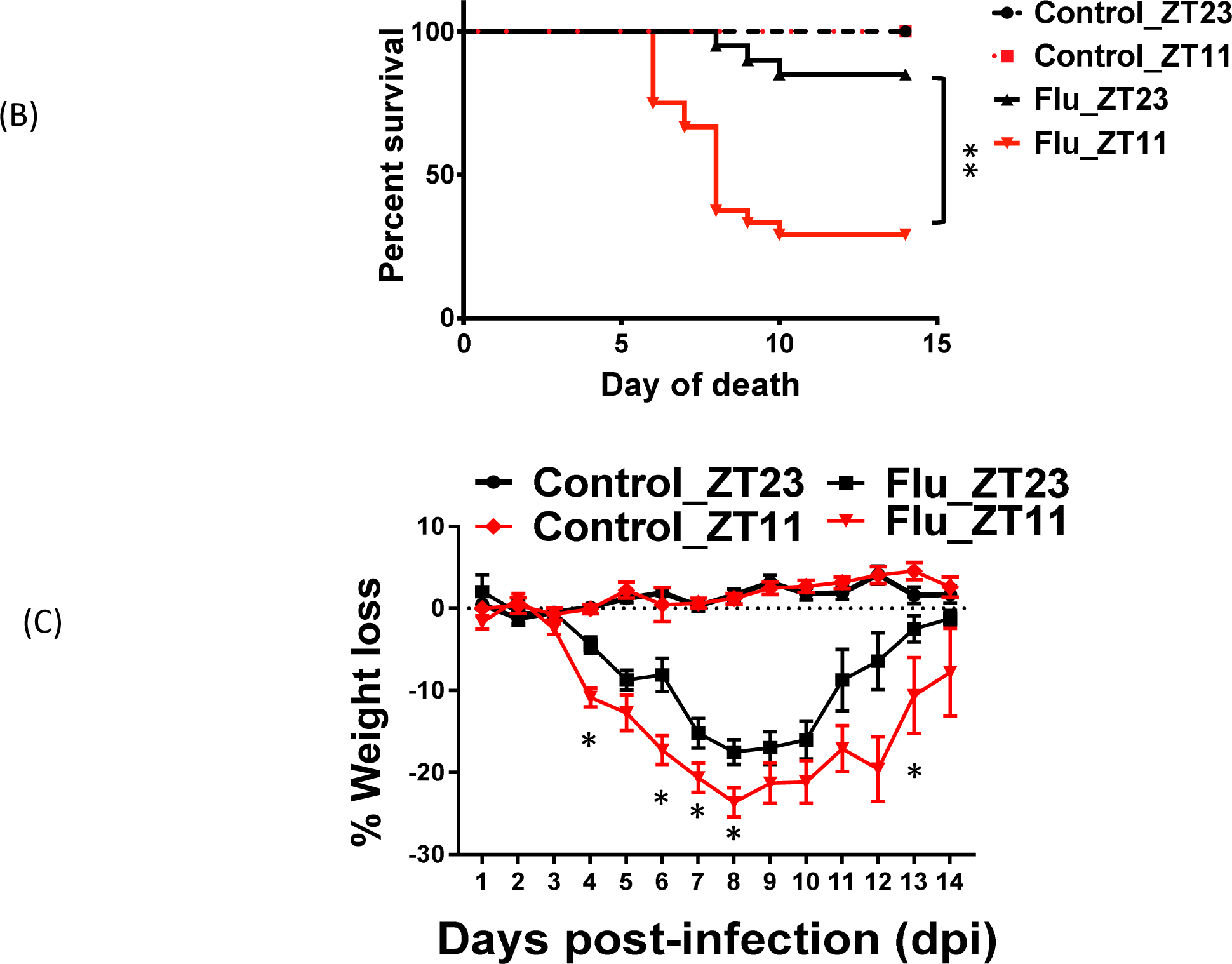
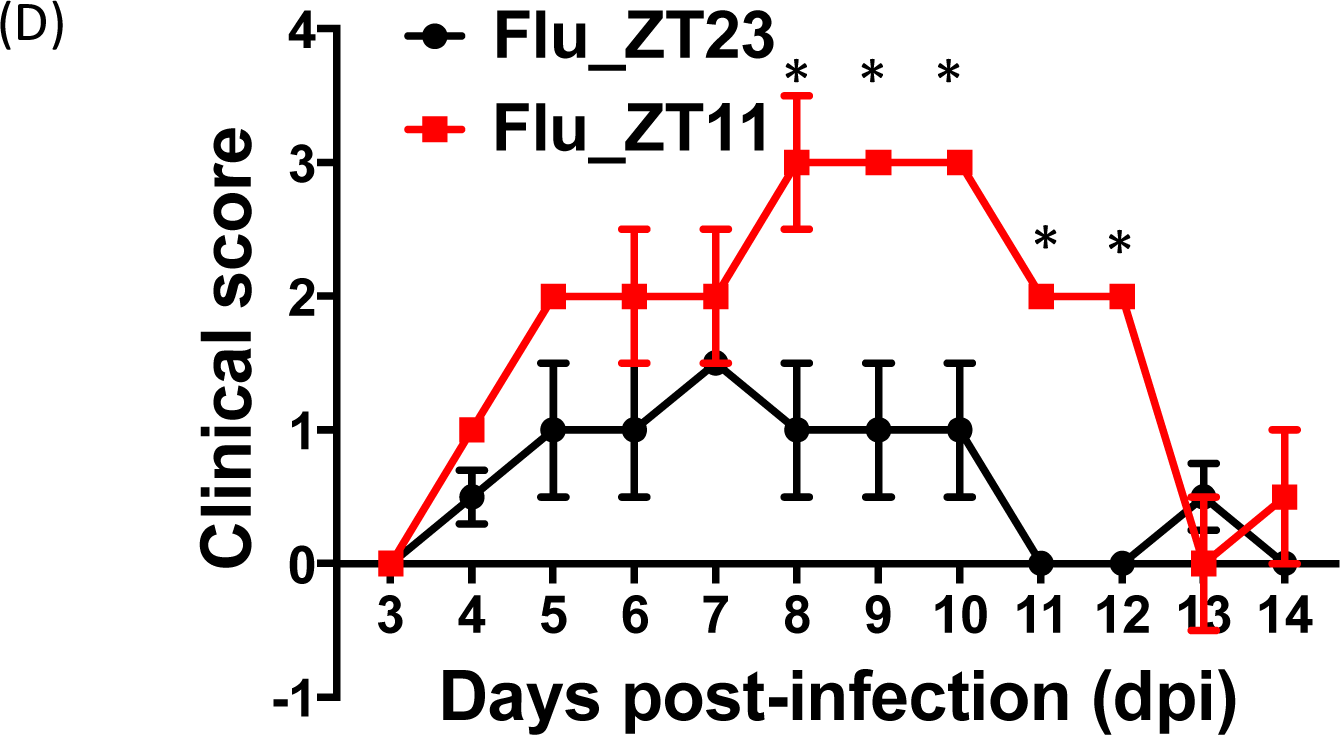
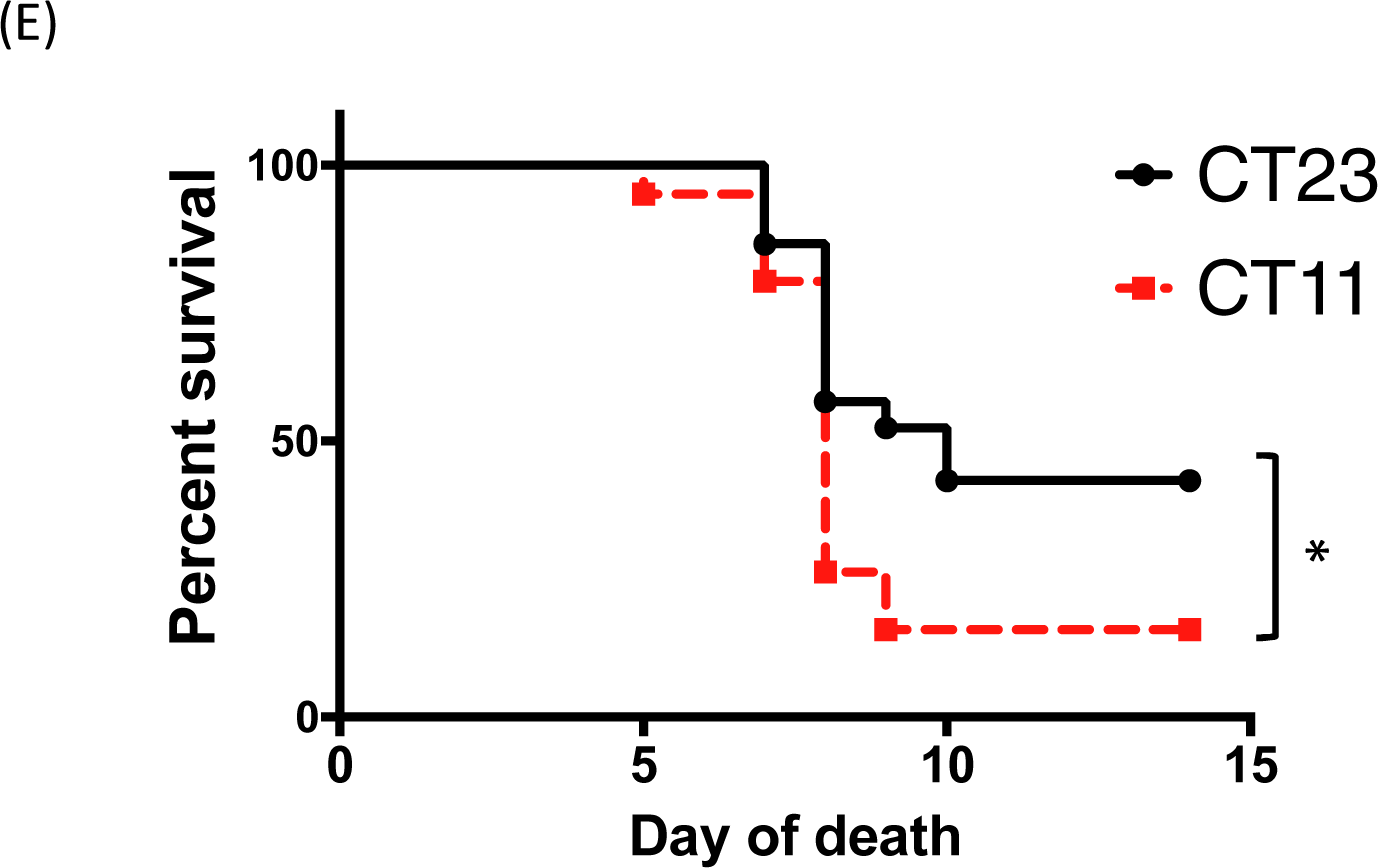
Time of day of infection affects survival following influenza A virus (IAV) infection by intranasal (i.n.) administration. *(A)* Experimental design: Two groups of mice were maintained in 12 hrs Light: Dark cycles. One group of mice was infected with 0.5LD_50_ at the start of the light cycle (ZT23), and for the other group i.n. IAV was administered at the start of the dark cycle (ZT11). *(B)* Survival curves are a composite of 3 independent experiments [total n = 17 per group, log-rank (Mantel–Cox) test, p < 0.0001]. *(C)* Disease progression is expressed as the percent of weight change following IAV infection. Compiled data are expressed as mean ± SEM. *(D)* Disease progression was also measured as clinical scores. Data represented as median ± SEM (total n = 17 per group; Student’s t test, *p < 0.05, **p < 0.01, ***p < 0.001). *(E)* Two groups of mice were maintained in constant darkness for 72hrs. One group of mice was infected with 0.5LD_50_ at the start of the light cycle (CT23), and for the other group i.n. IAV was administered at the start of dark cycle (CT11). Survival curves [total n = 8-12 per group, log-rank (Mantel–Cox) test, * p < 0.05].

These data are consistent with the hypothesis that the susceptibility to IAV induced mortality and morbidity depends on the time of day at infection.

### The temporal gating of IAV induced disease is circadian

To show that the diurnal differences in mortality and morbidity were due to endogenous circadian regulation as opposed to the effect of light:dark cycles, we repeated the above experiment in mice housed in constant darkness. The mice were infected with IAV at either CT23 or CT11 (which corresponds to the beginning of the rest phase and of the active phases respectively) with the same dose of IAV as above. We found that mice infected at CT11 had significantly higher mortality than the mice infected at CT23 (Fig. 1E), consistent with endogenous circadian rhythms controlling the outcome of the IAV infection.

### The time of day at infection affects viral clearance but not viral replication

To test if the difference in the outcomes were driven by a varying rate of viral replication, we measured viral titers in the lungs on days 2, 4, 6, 8 p.i. The titers were not different in the two groups at the earlier time points (Fig. 2A) when viral replication is known to peak (days 2,4 and 6). By day 8 p.i., more mice infected at ZT23 had cleared the virus than those infected at ZT11. Therefore, it is unlikely that the differences in mortality and weight trajectories can be attributed to viral replication, because clearance follows rather than precedes the mortality and morbidity observed. Thus, despite inciting higher inflammation in the ZT11 group, viral clearance is delayed. Further, since several previous studies have reported higher morbidity and mortality in females(13, 14), we also stratified the experiment by gender but observed no difference in viral kinetics. (Fig. 2B).

**Fig. 2:**
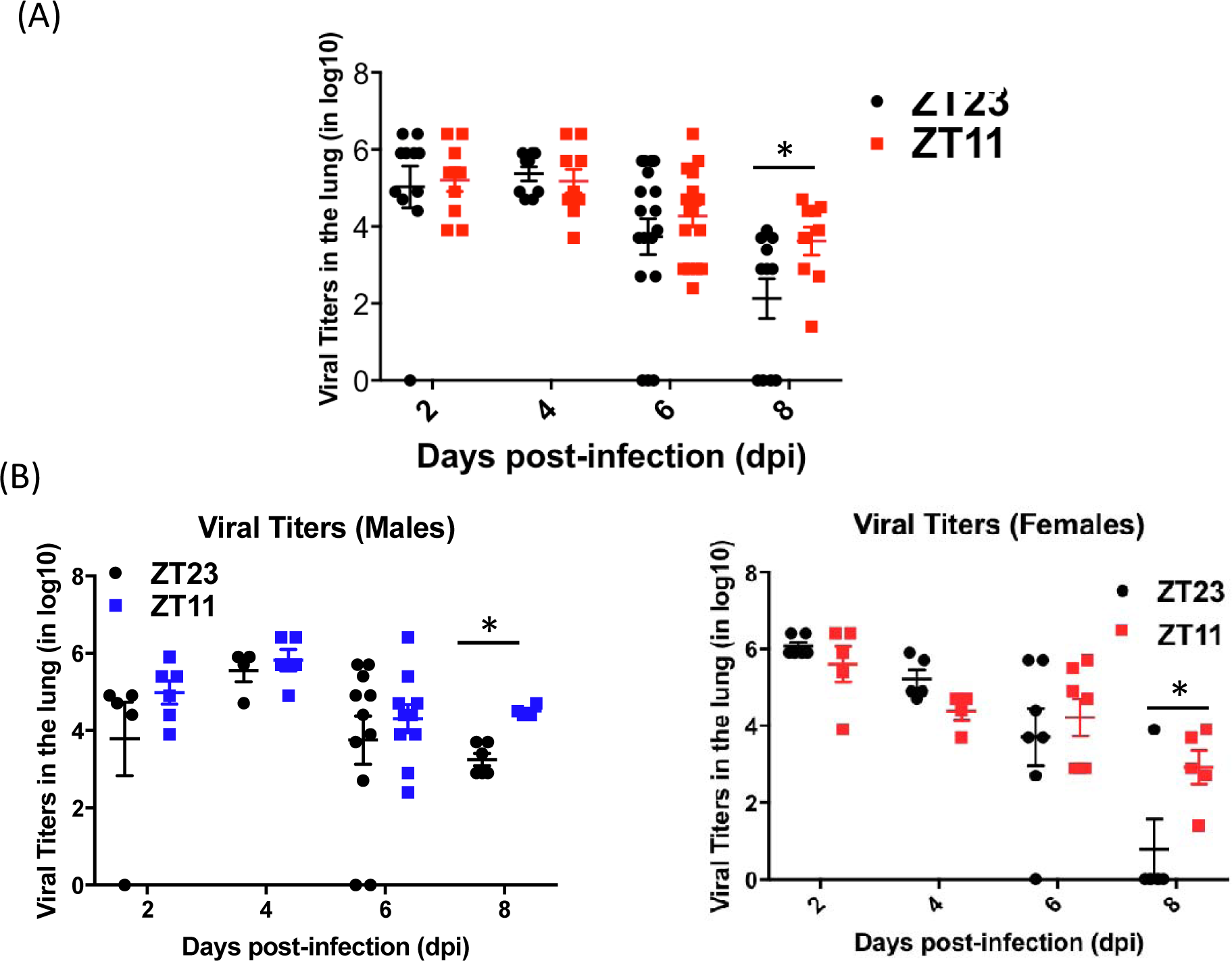
The time of day at infection affects viral clearance. After infecting mice at ZT23 or ZT11, viral titers were determined in the lung on 2, 4, 6, 8 (days post-infection) d.p.i. *(A)* Combined data. (n = 8-12 mice per group, student t test; *p < 0.05, **p < 0.01, ZT23 vs. ZT11). *(B)* Stratified by gender.

### Circadian gating of IAV-induced mortality and morbidity is associated with lung inflammation

Since there were no differences in the pulmonary viral loads between the groups infected at ZT11 and ZT23, we compared the extent of inflammation in these two groups. Mice infected at ZT11 had higher total BAL cell counts as early as 2 p.i. but this persisted even on day 6 (Fig. 3A & D). Further, we also collected BAL at 4-8hr intervals across 24 hrs from day 2 to day 3 p.i.. The total BAL cell counts were higher in the ZT11 group across all time points, proving that infection at ZT11 promoted more inflammation and this was not a function of time of dissection (Fig. 3A). However, there were no significant differences in the differential counts between the two time points (Fig. 3A & C). At baseline however, the BAL cell count did not oscillate across time (Fig. S2), although circadian oscillations in CD45^+^ cells have been reported in dissociated whole lung preperations(15).

**Fig. 3:**
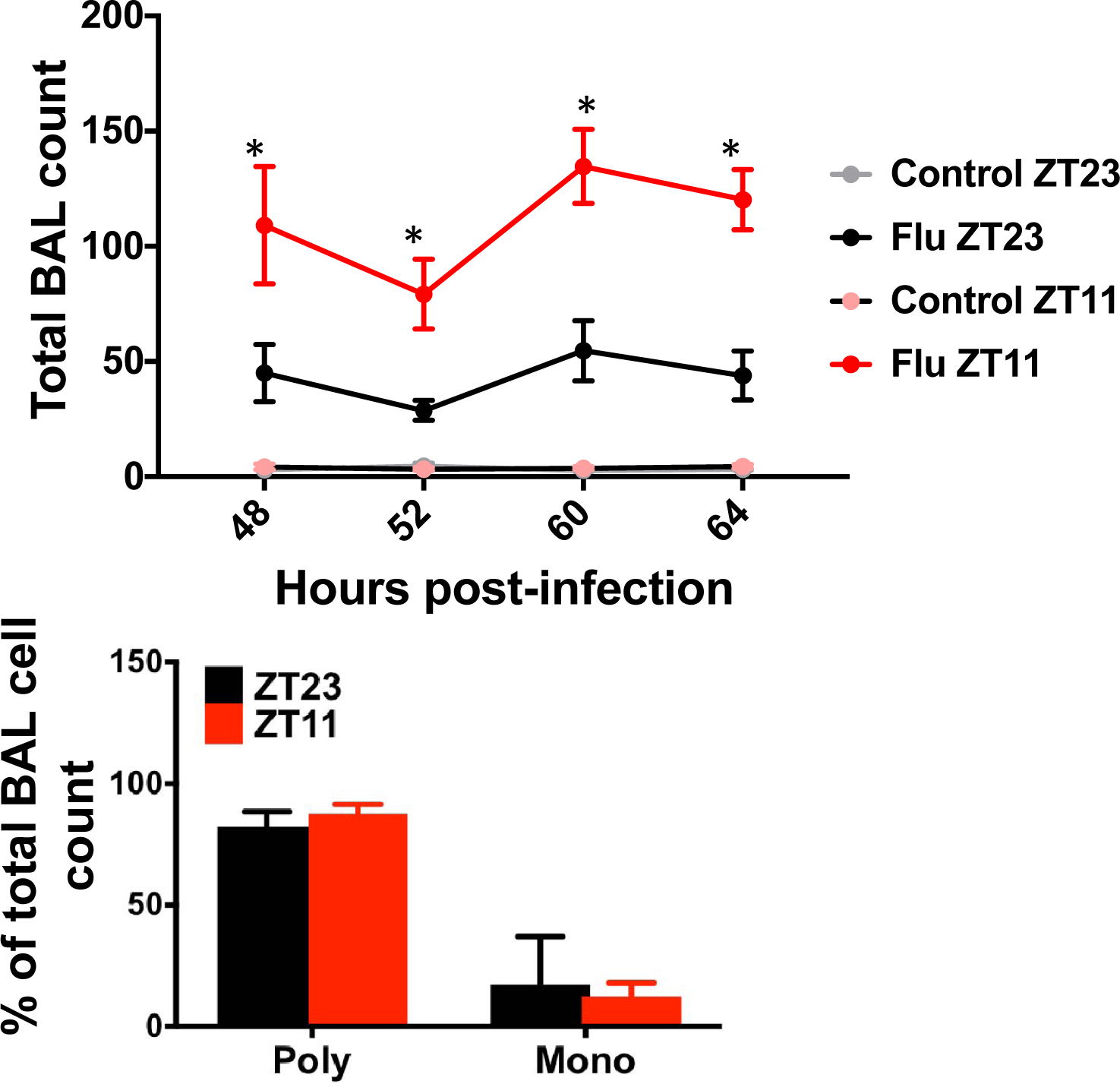
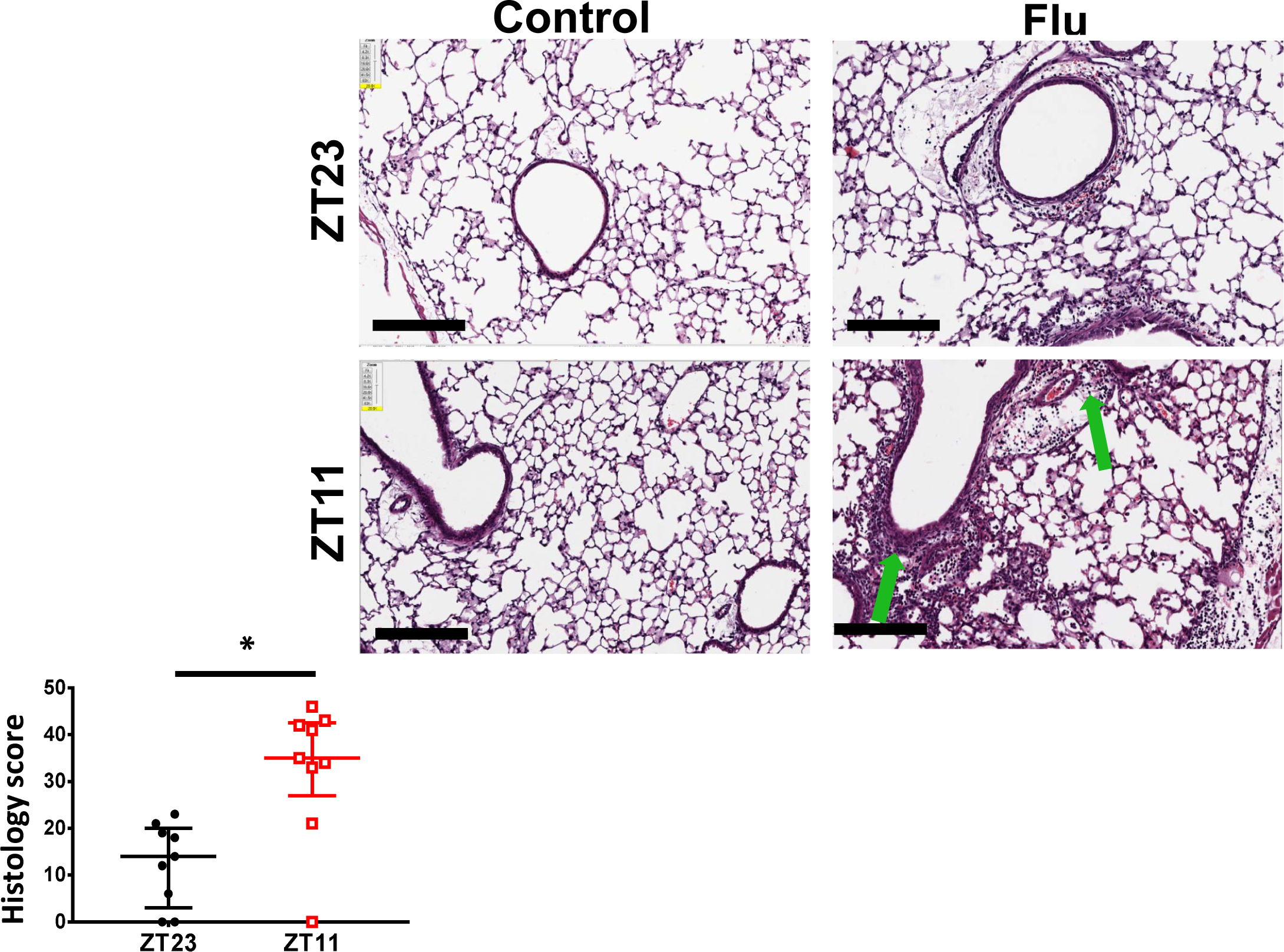
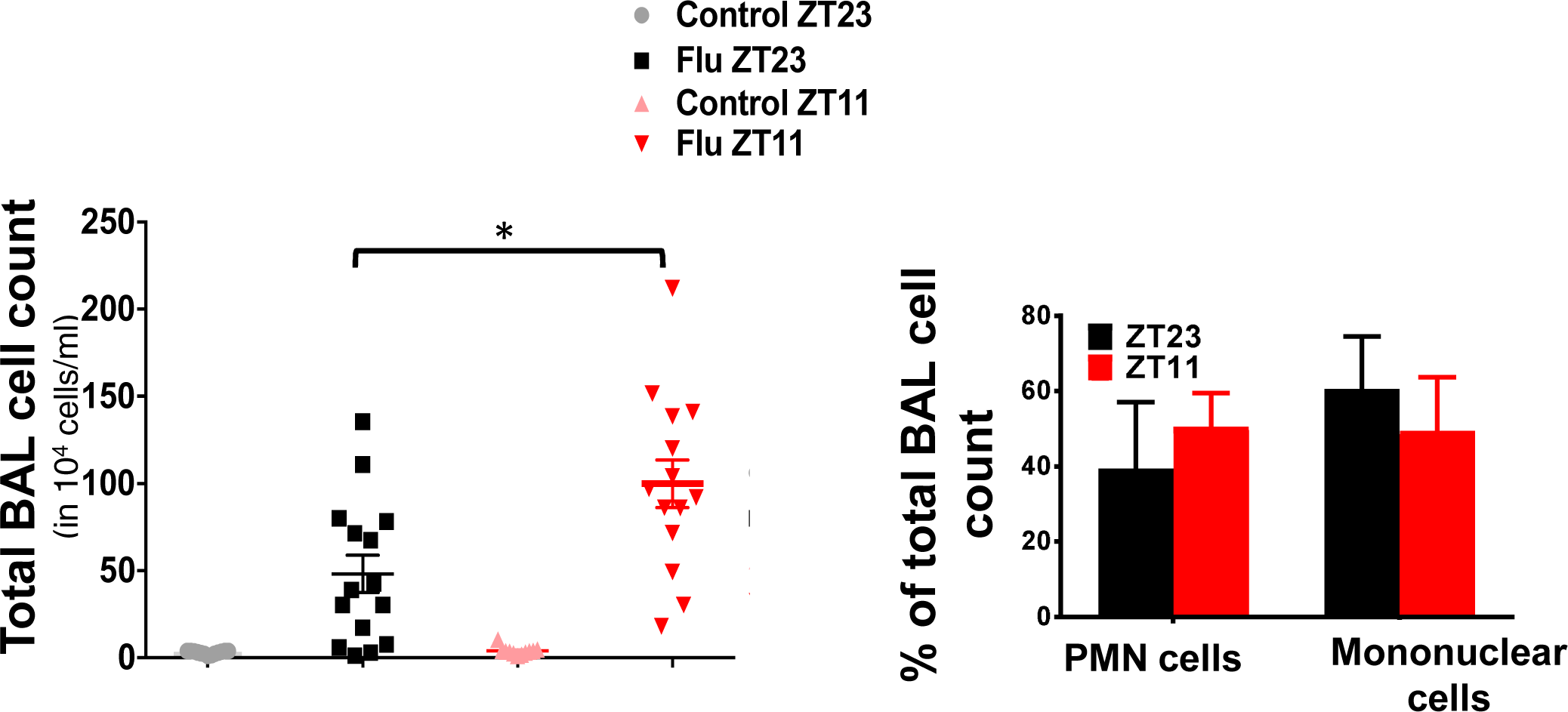
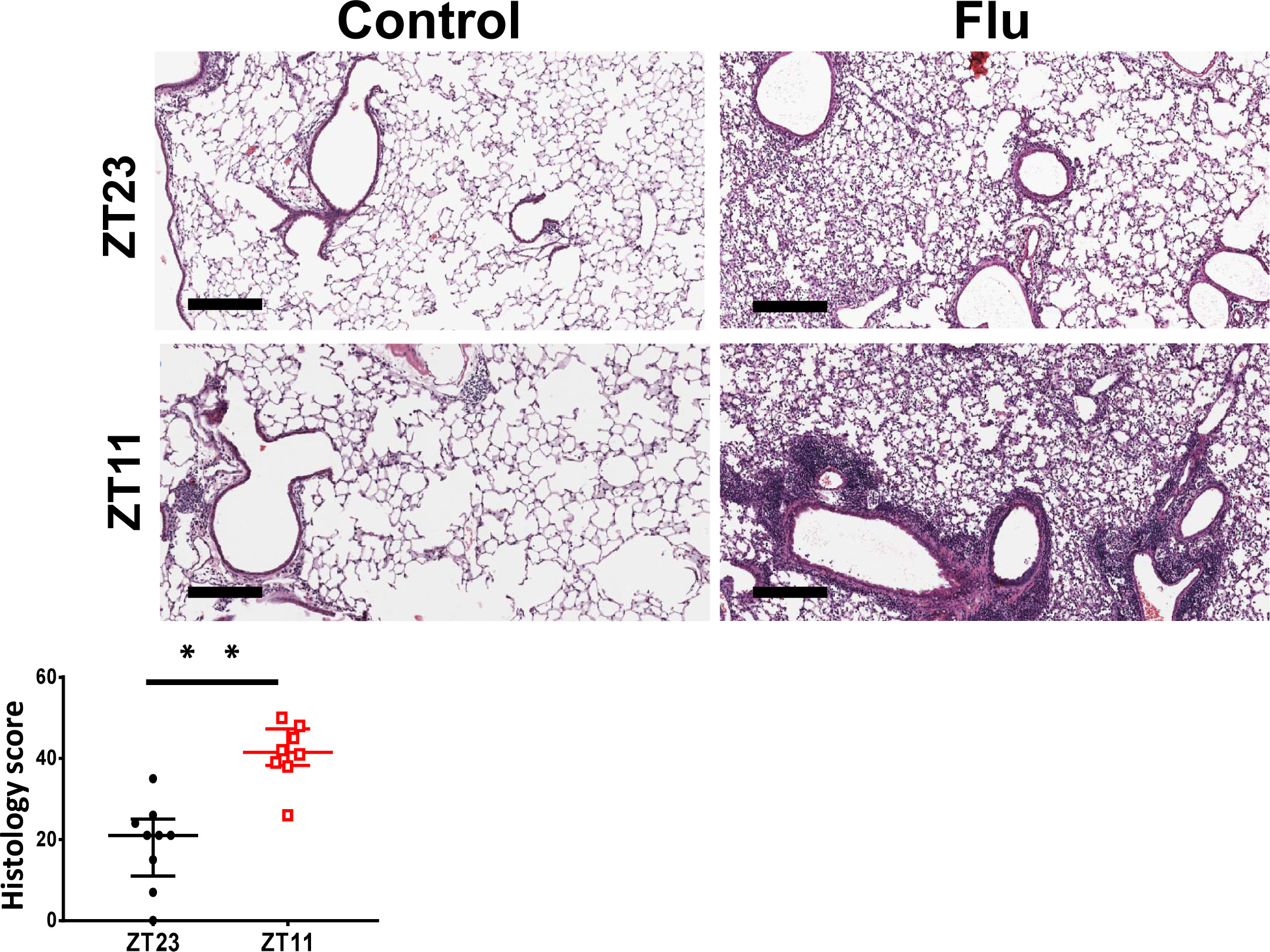
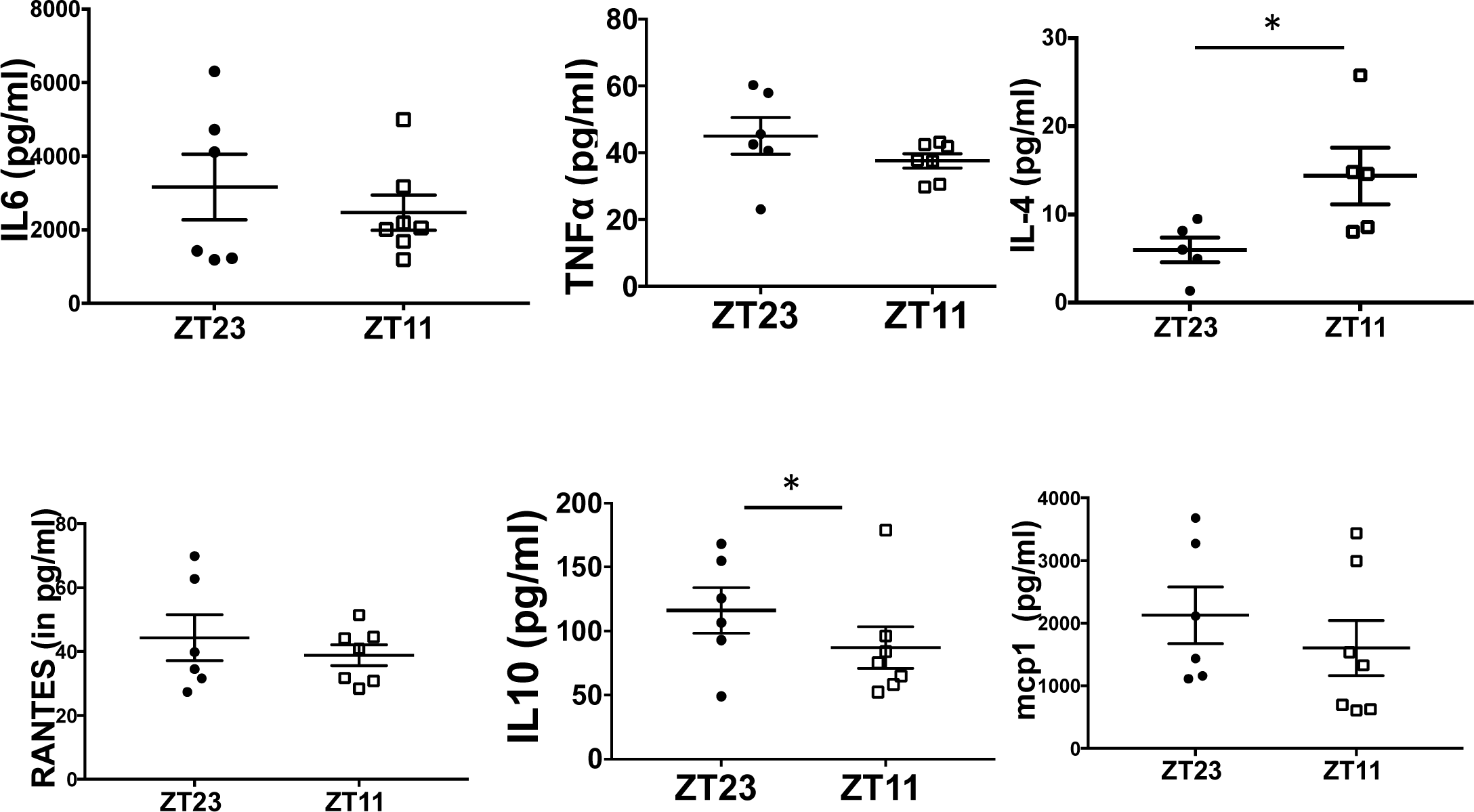
Temporal gating of IAV-induced mortality and morbidity is associated with lung inflammation. *(A)* Broncho alveolar lavages (BAL) were collected at the indicated time points post-infection from mice that received either IAV or PBS either at ZT23 and ZT11. Top Panel: Total BAL cell count 48-72 hours p.i. *Lower panel:* Differential of the cells from 48 hrs post-infection for both IAV infected groups. Data were compiled from three independent experiments and expressed as mean ± SEM (total n = 5-8 per time point, two-way ANOVA; *p < 0.05 for time of infection, NS for time of dissection). *(B) Top Panel:* Representative micrographs of H&E stained lung sections 2 days after sham (PBS) or IAV (50 PFU) treatment. (photomicrograph bar=200µm). *Lower Panel:* Severity of lung injury quantified using an objective histopathological scoring system. ((total n = 5-6 per group, student t test; *p < 0.05, ZT23 vs. ZT11) *(C) Right Panel:* Total BAL cell count on day 6 p.i. from mice who received either IAV or PBS at either ZT23 and ZT11. Data compiled from three independent experiments are expressed as mean ± SEM (total n = 9-15 per time point, two-way ANOVA; *p < 0.05, ZT23 vs. ZT11). *Left Panel:* Differential of the BAL cells from both IAV infected groups. *(D) Top Panel:* Representative micrographs of H&E stained lung sections 6 days after sham (PBS) or IAV (40 PFU) treatment. (photomicrograph bar=200µm). *Lower Panel:* Severity of lung injury quantified using an objective histopathological scoring system. ((total n = 7-9mice/ group, student t test; **p < 0.01, ZT23 vs. ZT11) *(E)* Cytokine levels in BAL. (n=6/group. student t test; *p < 0.05)

Mice infected at ZT11 had more lung injury on both days 2 and 6 p.i. (Fig.3B & 3D) based on more peri-bronchial inflammation, peri-vascular inflammation, inflammatory alveolar exudates and epithelial necrosis. Similar effects were also seen with X31 strain of influenza virus, suggesting that these effects are not restricted to one strain but rather reflect the circadian control of influenza infection overall (Fig S3). We assayed for a cytokine panel in the BAL on day 6 that correlates with peak viral load in the lung. There were no significant differences in the levels of cytokines Il6, Tnfα, Ccl5 or Ccl2 (Fig S4, 3E). However, the levels of Il4 were higher in the group infected at ZT11 than those infected at ZT23 (Fig. 3E). Il4 is associated with the development of type-2 adaptive immunity which is suboptimal for viral clearance in other respiratory viruses, such as rhinovirus(16). Thus, higher Il4 in the ZT11 group would be expected with the worse outcomes observed. IL10, a cytokine that suppresses inflammation in acute influenza(17), was higher in ZT23 group than ZT11. We speculate that the cytokines may peak later; we were not able to detect difference in other cytokines at this point. Interestingly, when we challenged mice with i.n. polyinosinic:polycytidylic acid (Poly I:C), a TLR3 ligand, at either ZT23 or ZT11, the results were reversed (Fig. S5), with mice infected at ZT23 exhibiting worse lung injury and higher total BAL cell counts. This finding suggests that the difference in outcomes in mice injected with IAV at different time points is not likely due to pathways downstream of TLR3. Considered together, our results are consistent with the hypothesis that endogenous circadian rhythms determine mortality and pathology in IAV infection by modulating the inflammatory response, rather than through an impact on viral load in the lung.

### The circadian gating of IAV infection is associated with a distinct transcriptomic signature

To determine whether the circadian control of IAV infection might be regulated at the transcriptomic level, we performed RNA sequencing of the whole lungs from mice infected with either IAV or PBS at ZT23 or ZT11 on day 6 p.i.. We chose this time point for transcriptomic analysis because the weight loss trajectories had clearly diverged but the levels of cytokines were not significantly different between the two groups. This would allow us to identify pathways that might explain the difference in outcomes.

Of a total 29,114 genes, 4667 had a ≥ 2-fold difference between the two IAV-infected groups. Although our experimental design was not set up to detect circadian rhythms in gene expression, 184 genes had a ≥ 2-fold difference from the control groups at ZT23 and ZT11 (Fig. 4A). Mice infected at ZT11 had a distinct transcriptomic profile compared to all other groups (Fig. 4B). As expected, innate immune pathways involving cell migration and pattern recognition are among the most enriched and are consistent with innate immunity-mediated circadian gating of the flu infection (*Table 1*). Further pathways enriched in the transcriptomic analyses, included various aspects of innate immunity and the “Role of hypercytokinemia/hyperchemokinemia in pathogenesis of Influenza”. Consistent with our other results and the literature (18, 19), the transcriptomic analyses provide further evidence that a state of hyper-inflammation is induced by infection at ZT11, to which the host eventually succumbs (Fig. 4B & 4C).

**Fig. 4:**
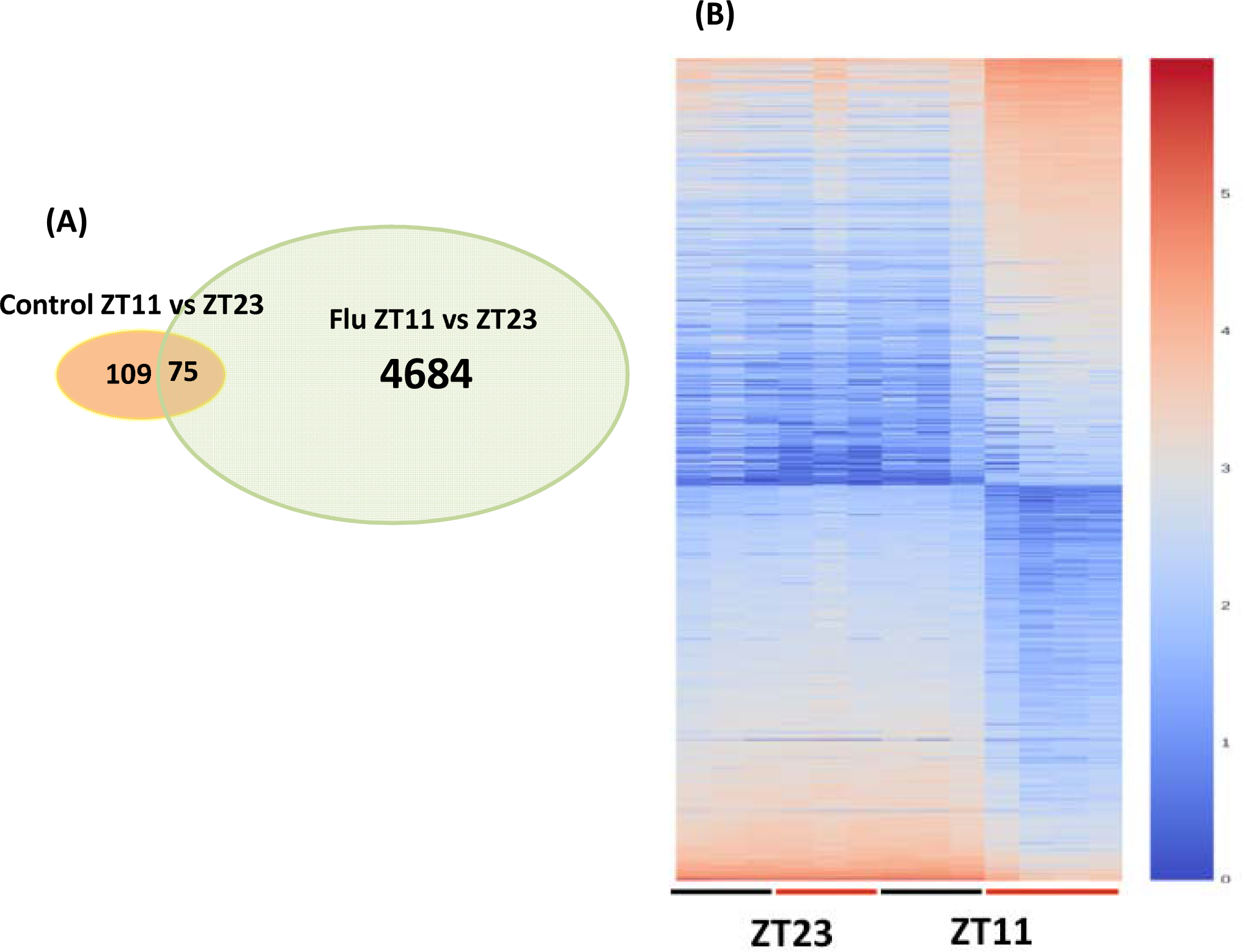
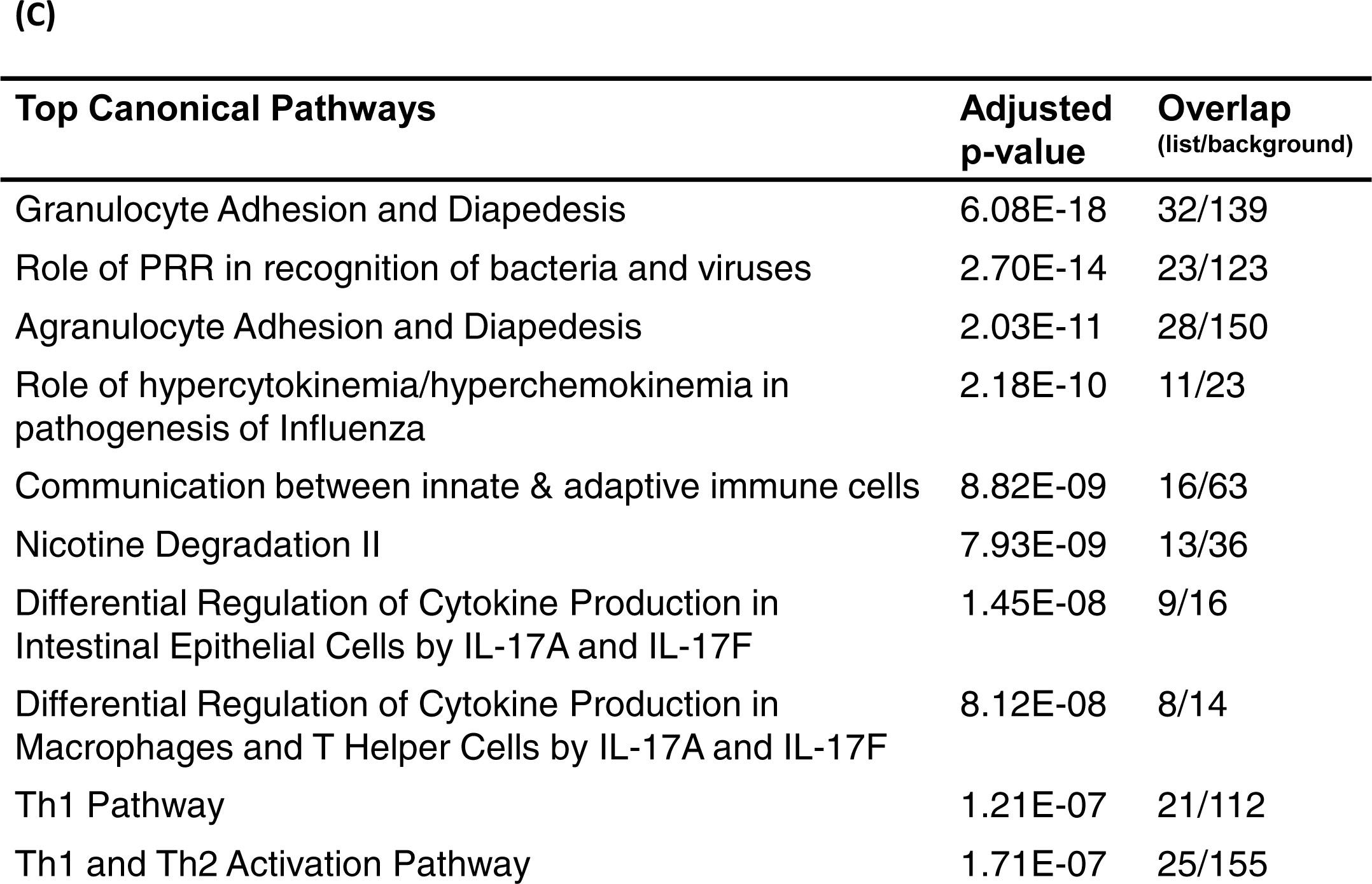
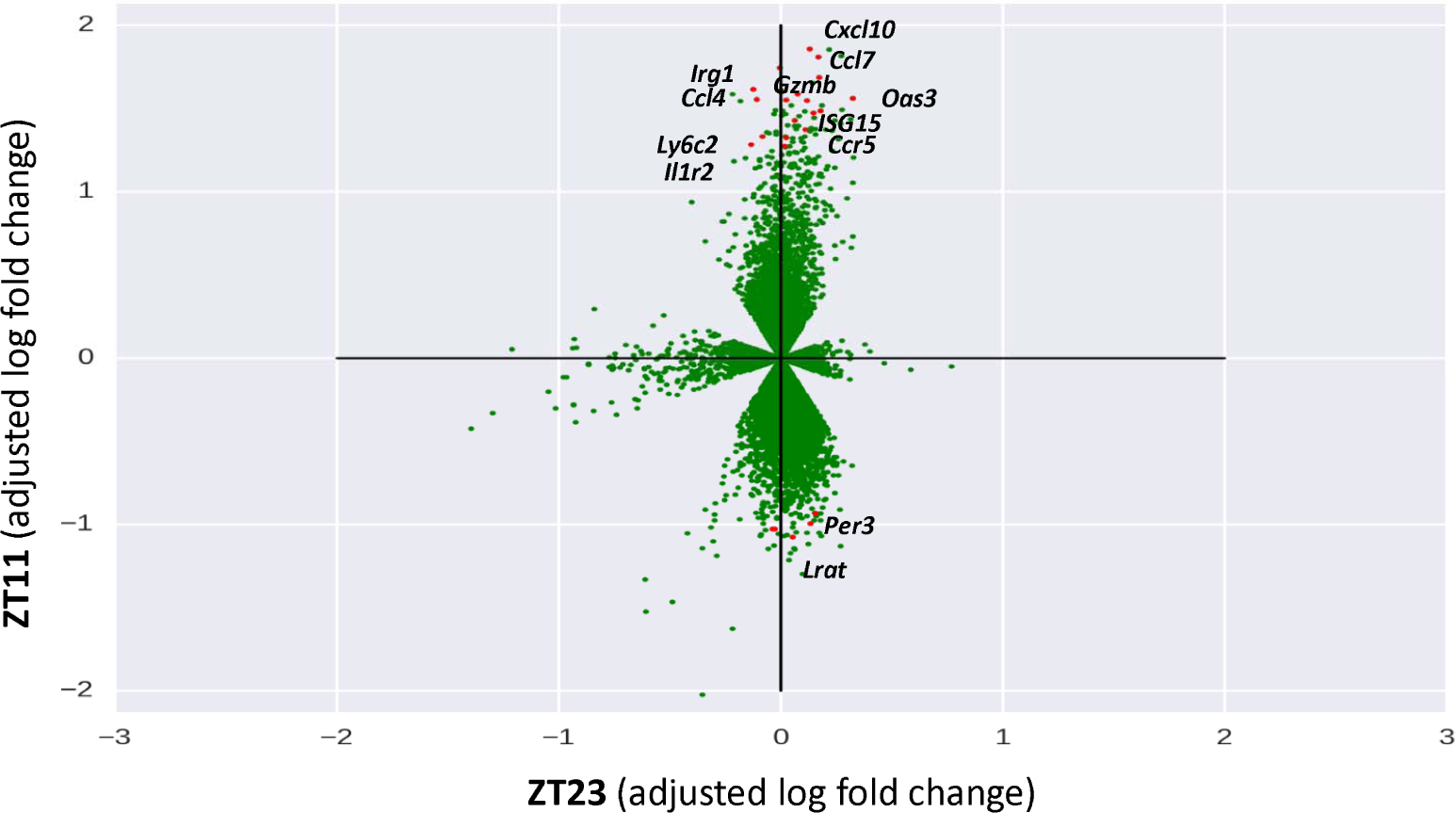
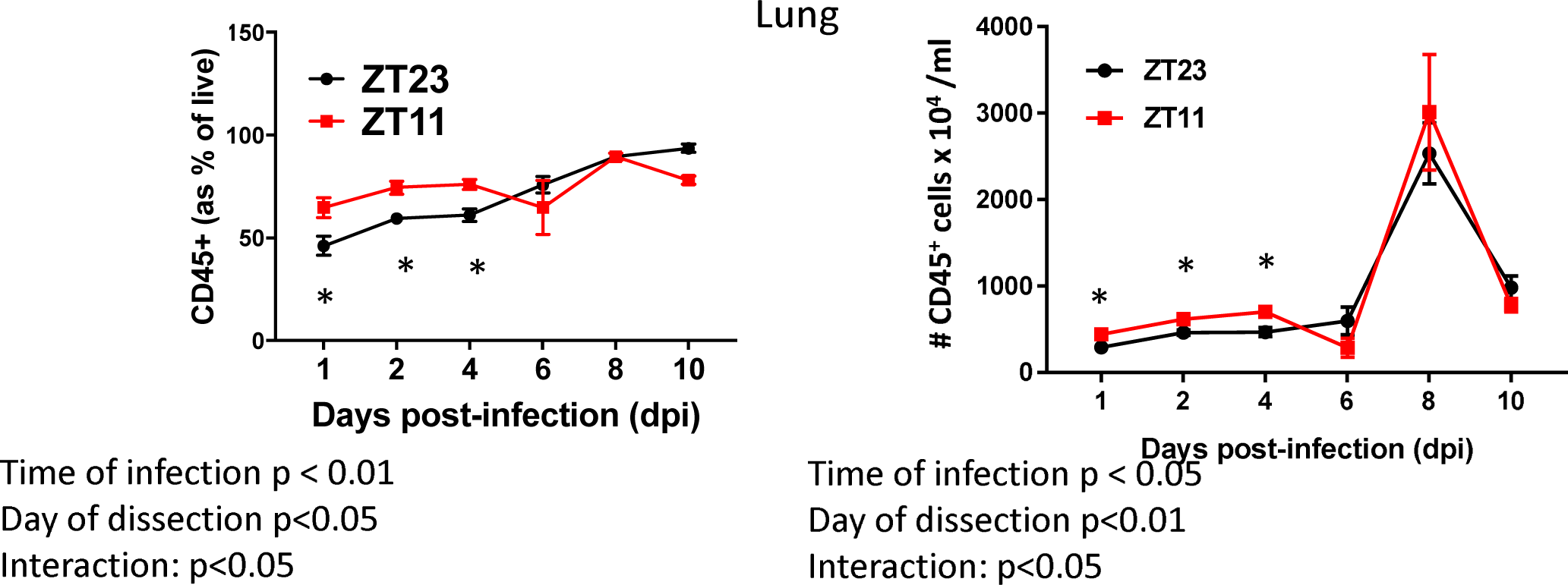
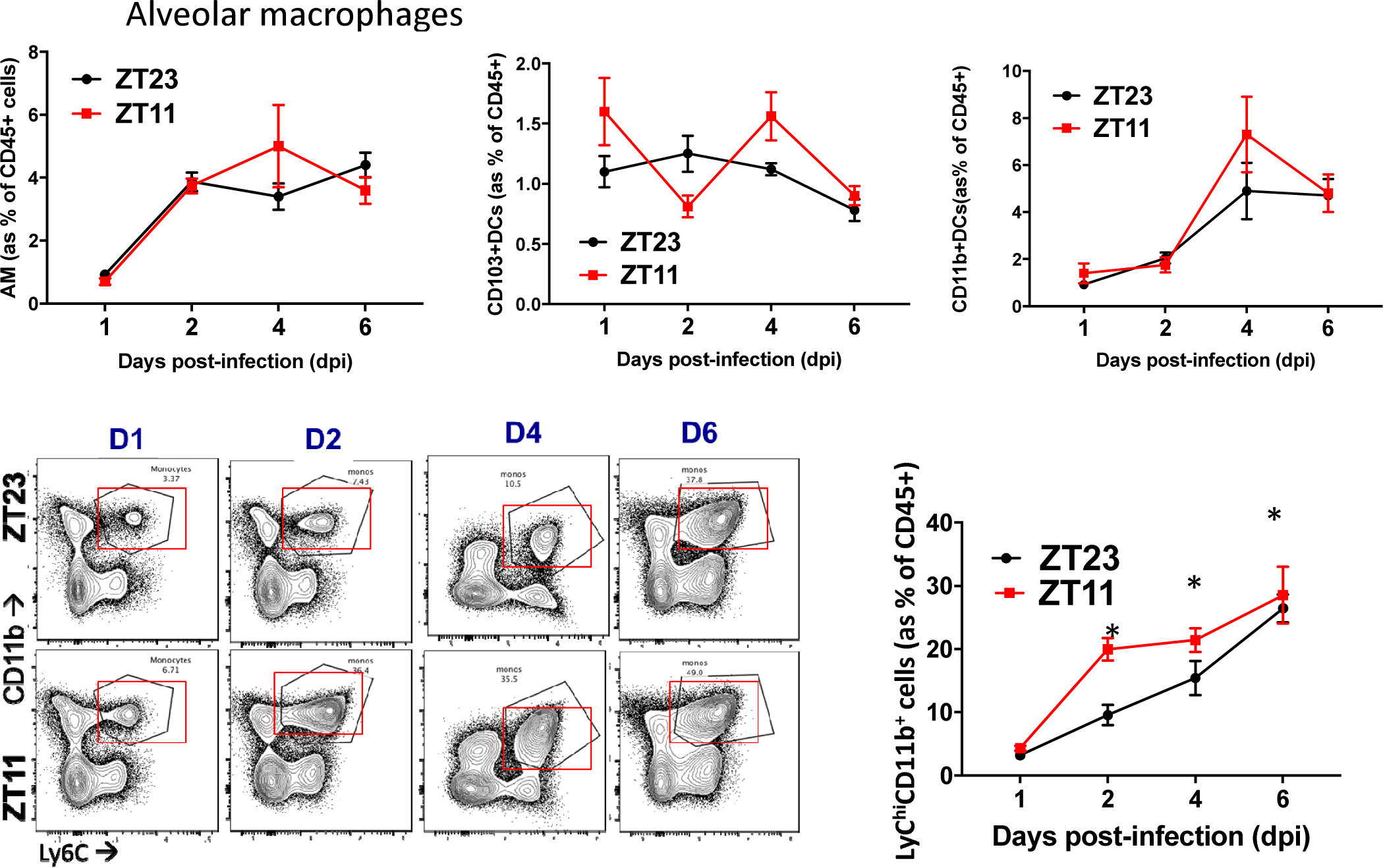
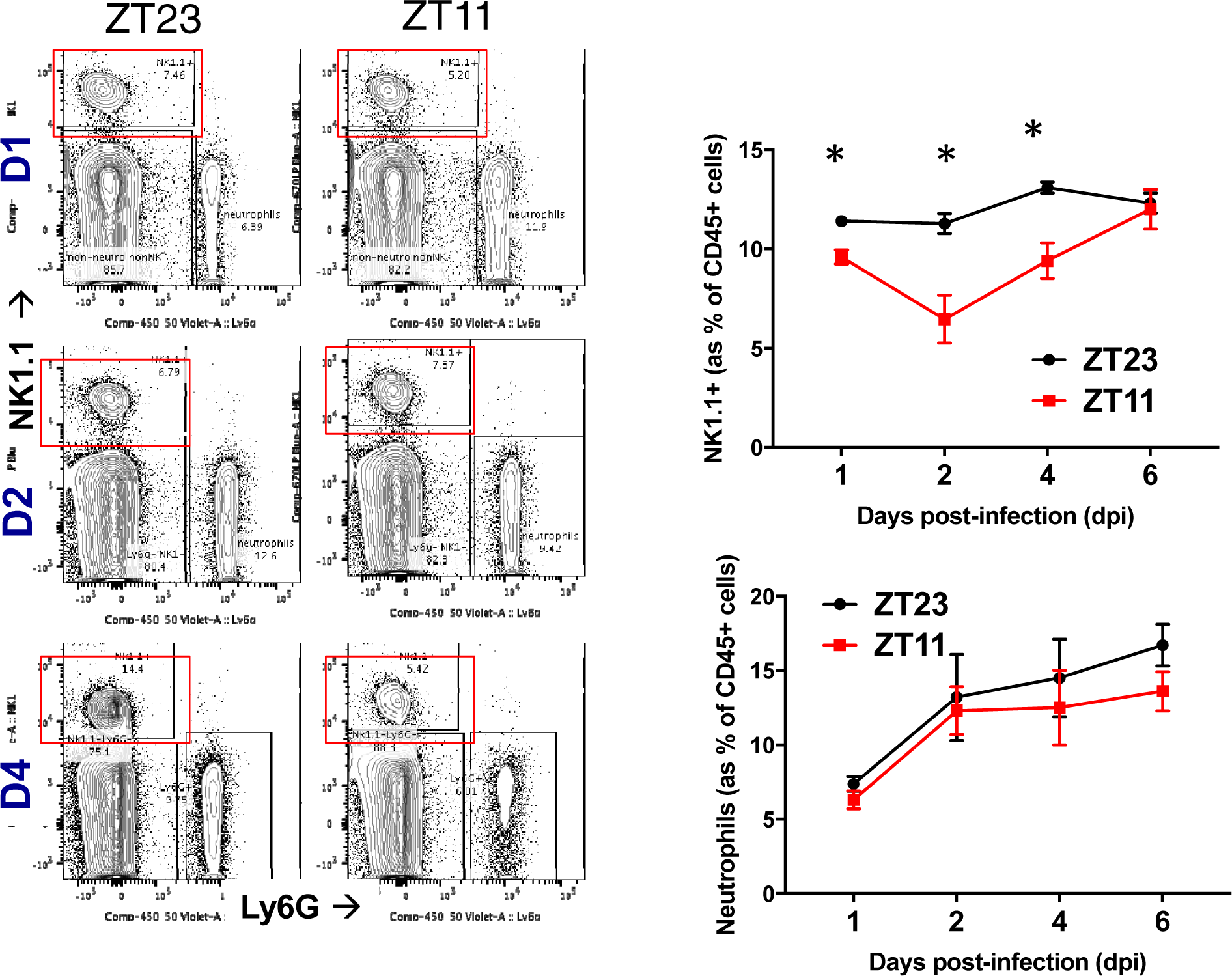
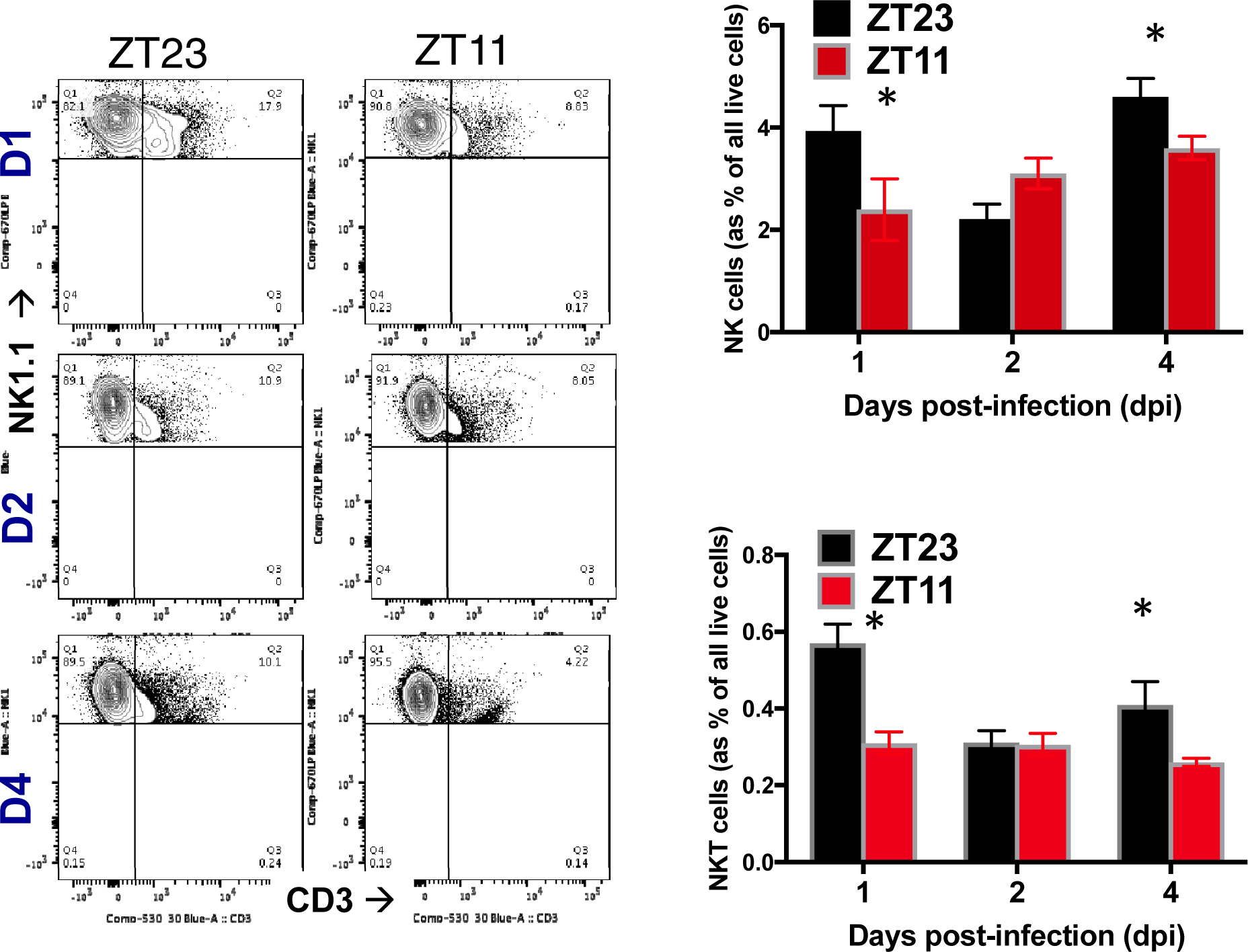
RNA samples from animals infected at ZT23 or ZT11, with either PBS or IAV, 6 days after infection were collected at ZT23 and ZT11 and used for RNA-Seq. *(A)* Venn diagram (sizes not to scale) depicting the number of differentially expressed genes. *(B)* Heatmap of the top 900 differentially expressed genes. *(C)* Plot of log adjusted fold change for ZT11 and ZT23, showing directionality of the most differentially expressed genes. Flow cytometry-based enumeration of the different innate immune cell populations in dissociated lungs following IAV infection at either ZT23 or ZT11. *(D)* CD45+ cells (as a % of live) and CD45+ cell numbers. 2-way ANOVA, p<0.01 for time of infection<0.05 for day of dissection and p<0.05 for interaction. *(E)* Macrophages, two subsets of dendritic cells (CD11b+ and CD103+) and Ly6C^hi^ monocytes with images from representative experiment. For monocytes, 2-way ANOVA, p<0.0001 for time of infection, p<0.001 for day of dissection and N.S. for interaction. *(F)* Neutrophils and NK1.1+ cells. 2-way ANOVA, p<0.05 for time of infection<0.05 for day of dissection and p<0.05 for interaction. *(G)* NK cells and NKT cells. For NK cells 2-way ANOVA, p=ns for time of infection and p<0.01 for day of dissection and p<0.05 for interaction. For NKT cells 2-way ANOVA, p<0.05 for time of infection and N.S. for day of dissection or interaction. (N=4-7/group per time point. Two-way ANOVA for time of infection and time of dissection). Representative results from one experiment. Experiment repeated with similar results 3-5 times.

### NKT cells and NK cells are associated with the temporal gating on influenza infection

To identify which cell types mediate the circadian difference in the response to IAV, we determined the proportion of CD45+ populations in the lungs of mice infected at either ZT11 or ZT23 on days, 1, 2, 4, 6 and 8. First, similar to our results from BAL analyses, we found that mice infected at ZT11 had a higher total CD45+ cell count in the early phase of infection (days 1-4) (Fig. 4D). There were no significant differences in the numbers or percentage of macrophages, neutrophils or CD11b+ DCs in the lungs, between the two groups (Fig. 4E & 4F). The percentage of CD45+ NK1.1^+^LysG^-^cells was higher in the ZT23 group than the ZT11 on days 1-4 p.i.. While both NK cells (CD45^+^LysG^--^Nk1.1^+^CD3^-^) and NKT cells (CD45^+^LysG^--^Nk1.1^+^CD3^+^) were higher in the ZT23 group, the latter seemed more significantly associated with time of infection. (Fig. 4G; p<0.01 on 2-way ANOVA for time of infection). The proportion of CD45^+^SiglecF^-^Ly6G^-^CD11b^+^Ly6c^hi^ monocytes was higher in the ZT11 p.i. group, suggesting a state of heightened inflammation (Fig. 4E). While, some recent reports have implicated circadian regulation of the adaptive immune system in disease pathology(20) and lymphocyte trafficking(21), others have not(22). In our model, by day 8 or 10 p.i. there were no significant time dependent differences in the total CD8+ cells or activated CD8+ cells in either the lung or mediastinal LNs (Fig S7).

Thus, based on these findings, we propose that NK and NKT cells may protect the host from influenza related inflammation in the ZT23 group, while monocytes may predispose to the worsened inflammation in the ZT11 group.

## Discussion

Here we demonstrate that lung inflammation induced by IAV is modulated by circadian rhythms. We also show that circadian regulation of host tolerance and not viral kinetics determines the observed outcomes. This effect was independent of light:dark cycles. This temporal difference in outcomes based on time of inoculation is consistent with recent trials of vaccination that demonstrate that time of day affects antibody responses (12, 23, 24).

One of the most interesting findings from our study is the elucidation of the role of hyper-inflammatory cascades in mediating the circadian gating of IAV. We show that infection at ZT11 results in higher total BAL cell counts, worse lung pathology and greater number of total lung CD45+ cells – evidence for increased inflammation. Pathway analyses of whole lung transcriptomic data are also consistent with higher inflammation causing the disparity in outcomes between infection at ZT11 vs ZT23. Moreover, early in infection, when the weight loss and mortality trajectories have diverged in the ZT11 and ZT23 groups, there is no significant difference in viral titers. This contrasts with prior studies of other pathogens that attribute the cause of circadian variability in response to variation in abundance of pathogen or its direct replication in the host(9, 11). Gibbs et al, reported a difference in the bacterial load within 24-48 hrs of infection with *Streptococcus pneumonia*. More relevant to viral replication *in vivo*-Ehlers et al reported that antiviral activity tracked with viral nucleic acid abundance in Sendai virus infection in *Bmal1^-/-^*mice where the clock is disrupted. Similarly, viral replication in the host was responsible for the circadian control of infection by a luciferase-expressing Murid Herpes strain(9) in mice and for influenza in *Bmal1^-/-^* fibroblasts.(25)

We determined viral load by infectivity assay(26), which is directly determines viral activity. This is preferable to the determination of viral loads through viral RNA expression, which may be confounded by other factors, such as the presence of defective viral genomes (DVG). Overall, our results are consistent with existing reports that mortality in influenza infection may be caused the extensive activation of immune pathways(27), rather than just viremia or extra-pulmonary effects(27-29).

The temporal gating of the host’s response to influenza infection is also supported by some recent trials where the antibody response correlated with the time of vaccination(12, 23). IAV, in turn, seems may influence host circadian rhythms (30, 31).

In our study, NK and NKT cells were higher in the ZT23 group than in the ZT11 group, early in the infection’s course. NK cells have direct cytolytic activity towards virally infected cells and accumulate at the site of infection, peaking around days 4-5 p.i. which is consistent with our results. Mice lacking *NCR1*, the predominant activating NK cell receptor succumbed to IAV early on, supporting the protective role of activating NK receptors in IAV(32). Further, NK cells and cytolytic function has been shown to be under circadian control in both humans(33) and animal models(34-36). In splenocytes obtained from *Per1^-/-^* mutant mice the rhythms of cytolytic activity, cytokines and cytolytic factors (such as granzyme B and perforin) and gene expression was significantly altered(35, 36). In the lungs of rats exposed to chronic jet lag, NK cytolytic activity was suppressed and tumor growth was stimulated(34). Thus, the existing literature is consistent with a role for NK cells in the circadian regulation of immune system, although not much is known about NKT cells in this context. However, given that the expression of *gzmb* and *prf1* were not significantly different between the two groups [Fig S8], we speculate that in our model NK cells may be acting primarily through non-cytolytic or immunosuppressive pathways. The second possibility is that the augmented balance between NK cells and Ly6C^hi^ monocytes in the ZT23 group may favor a milieu conducive to efficient viral clearance and a well-contained state of inflammation. Our results are consistent with other reports where monocytes have been shown to induce excessive inflammation by different agents (Listeria and VSV) in other organs(3, 10). In one model, the core clock gene *Bmal1* induced Ccl2(3) while in another *Rev-erb*α suppressed Ccl2(10, 37). The exact mechanisms underlying the role of both NK and NKT cells in mediating circadian regulation of lung inflammation may be specific to influenza. This remains to be elucidated.

Several studies aiming to link the circadian clock with inflammation use disruption of the non-redundant core clock gene, *Bmal1* (5, 9, 11, 30, 31, 38, 39) to implicate the molecular clock. However, *Bmal1* has a wide range of “non-circadian” effects(40). We suspect that some of these off-target effects specific to *Bmal1* may have resulted in the differences between our conclusions and those of other reports. Although weight loss was more profound in both models of *Bmal1* deficiency (vs WT), but not in *Per2^-/-^*mice, the effect was larger in magnitude in the embryonic *Bmal1^-/-^*mice than in the adult-inducible model *(Bmal1^fl/fl^ER^cre^)* supporting a confounding role of non-circadian functions of *Bmal1(11)*.

There is an abnormally high neutrophilic response in response to LPS mediated by Cxcl5, in mice where *Bmal1* was deleted specifically in the ciliated club cells of the lung (*CCSP-Bmal1)*. However, in response to Listeria, *Nyugen et al(3)* noted a preponderance of inflammatory Ly6c^hi^ monocytes in the spleen of *Bmal1l^Loxp/LoxP^Lyz2^Cre^* mice. More recently, Poullaird et al, noted increased neutrophils in response to LPS in *Rev-erb*α*-/-*mice at ZT0 but not at ZT4(41). While it is probable that neutrophils may serve as the effector cells for the circadian regulation of TLR4 signaling pathways(39, 41, 42) this mechanism does not underlie the response to viruses. In fact, in a global *Bmal1^-/-^*mice, there was no significant differences between the cellular responses to Sendai Virus(11).In other studies, the effect of *Bmal1* on viral nucleic acid expression(9) or mortality(30) was described, but specific cellular mediators in the host’s immune repertoire were not examined. In mechanisms specific to circadian regulation of viral infections, oscillations of the pattern recognition receptor, TLR9 has been reported(43).

In conclusion, our work demonstrates that time dependent regulation of influenza infection and its consequences are mediated by circadian regulation of host tolerance pathways and not directly through viral replication. Most therapeutic strategies for influenza are directed towards elimination of the virus. Targeting host tolerance via circadian-controlled pathways, may provide novel therapeutic possibilities. Although the circadian system may regulate the host response to many pathogens, it seems that the nature of the interaction varies with each dyad. Specific elucidation of how each infectious agent intersects with the host circadian machinery may be necessary to uncover therapeutic opportunities to modulate the relevant immune response.

## Methods

### Mice, Virus and Infection

Specific pathogen-free 8-12 week old C57bl/6J mice were purchased from Jackson lab. For influenza infections, mice were lightly anesthetized with isoflurane and infected intranasally (i.n.) with 30-70 PFU (based on body weight; LD50 is 100PFU) of PR8 strain or 10^4^ PFU of X31 strain of influenza virus, respectively, in a volume of 40 µl. For serial evaluations (for weights, scores, titers and cell counts) animals were assessed at the same interval from the time of infection. All animal studies were approved by the University of Pennsylvania animal care and use Committee and met the stipulations of the Guide for the care and Use of Laboratory animals. For experiments described in Fig 1 (A-C), animals were kept under 12hr LD cycles.

### Viral Titration

Lungs were harvested at different time points following infection, as indicated in the specific experiment. Lungs were extracted, homogenized in PBS-gelatin (0.1%), and frozen for preservation. The presence of influenza virus was evaluated using MCDK cells with 1:10 dilutions of the lung homogenates at 37°C. After 1 h of infections, 175 µl of media containing 2µg/ml trypsin was added and the cells were further incubated for 72h at 37°C. A total of 50 µl of medium was then removed from the plate and tested by hemagglutination of chicken red blood cells (RBCs) for the presence of virus particles. The hemagglutination of RBCs indicated the presence of the virus.

### Flow cytometry

Lungs were harvested after PBS perfusion through the right ventricle. The lungs were digested using DNAse II (Roche) and Liberase (Roche) at 37°C for 30 mins. Dissociated lung tissue was passed through a 70 um cell strainer, followed by centrifugation and RBC lysis. Cells were washed and re-suspended in PBS with 2% FBS. (Details of the antibodies in table S1). 2-3 × 10^6^ cells were blocked with 1ug of anti-CD16/32 antibody and were stained with indicated antibodies on ice for 20-30 minutes. No fixatives were used. Flowcytometric data was acquired using FACS Canto flow cytometer and analyzed using FlowJo software (Tree Star, Inc.). All cells were pre-gated on size as singlet live cells. All subsequent gating was on CD45+ in lung only. Neutrophils were identified as live, CD45^+^, Ly6G^+^ cells. Ly6C^hi^ monocytes were identified as live, CD45^+^Ly6G^-^Ly6C^hi^CD11b^+^ cells. NK cells were identified as CD45^+^Ly6G^-^LysC^-^NK1.1^+^ cells. In some experiments where indicated in the figure legend, an exclusion gate for neutrophils and T, cells (Ly6G, CD4 and CD8) was applied. Alveolar macrophages were identified as CD45^+^ Ly6G^-^Siglec F^+^; DCs were identified as live, CD45^+^Ly6G^-^SiglecF^-^CD11c^+^MHCII^+^ cells and further classified into CD103^+^ conventional DCs or CD11b^+^ DCs. Day 6 onwards, T cells were identified in mediastinal LNs as CD45+, either CD4+ or CD8+ cells. Activated cells were further differentiated as CD44^+^, CD62L^lo^. Gating strategy shown in Fig S5 and S6.

### Histology and BAL cytology

Flu infected and control mice were euthanized by CO2 asphyxiation and their tracheas cannulated with a 20 G flexible catheter (Surflo, Terumo, Philippines). The lungs were gently lavaged with 600ul of PBS in four passes. The supernatant from the first pass was collected and used for cytokine analyses. The cells from all four passes was pooled and re-suspended in 1 ml of PBS and counted using a Nexcelcom cell counter. Lungs were fixed by inflation with 10% buffered formalin at 20 mm H2O of pressure, paraffin embedded and stained with HE stain and PAS. Stained slides were digitally scanned at 63 × magnification using an Aperio CS-O slide scanner (Leica Biosystems, Chicago IL). Representative images were taken from scanned slides using Aperio ImageScope v12.2.2.5015 (Leica Biosystems, Chicago IL)

### BAL cytokine quantification

Customized panels of chemokine/cytokine were designed using MILLIplex Multiplex Assay and measured using Luminex (EMD Millipore) using the RIA core at the Institute for Diabetes, Obesity and Metabolism, University of Pennsylvania.

### Statistics

All statistical analyses were performed using STATA 11.0. Unpaired t-test or ANOVA was used for normally distributed data while Mann-Whitney as used for data without a normal distribution and for discrete scores (for lung histology).

### PCRs and Primers

#### RNAseq

RNA was extracted from whole lung homogenate using the RNA MiniElute kit as per the manufacturer’s protocol. QC was performed and only samples with RINs > 7 were used for sequencing. Library preparation was performed on 400 ng DNAse-treated RNA using the Illumina Truseq kit. Sequencing was done using HiSeq 2500 sequencer (Illumina) housed at BGI, CHOP to generate 2×100 strand specific paired-end reads. We obtained 30-50 mi pairs of reads per sample. Samples were aligned to a mouse reference genome on an in-house resampling-based normalization & quantification pipeline(44) and compared to existing gene annotations (ENSEMBL) to identify novel loci and isoforms. Differentially expressed genes were identified using a false discovery rate-based control for multiple testing. Finally, Ingenuity & GSEA were used to assess fully effects on key pathways and mediators.

#### Quantitative PCR

TaqMan gene-expression assays were used to measure mRNA levels for genes of interest. Eukaryotic 18S rRNA or GAPDH (both from Life Technologies) was used as an internal control. The samples were run on a LightCycler480 real-time PCR thermal cycler (Roche), and the relative ratio of the expression of each gene was calculated using the 2^δδCt^ method.

#### Transcriptomic Analysis

The RNA-Seq reads were aligned to the mouse genome mm10.GRCm38.p5 using STAR version 2.5.3a (Dobin et al., 2013). Following alignment, the normalization and quantification procedures were performed with the PORT version 0.8.2a-beta pipeline (**http://github.com/itmat/Normalization**). Gene level quantification was done by PORT with Ensemblv90 annotation. The goal of the transcriptomic analysis was to evaluate differential response to the IAV infection in the ZT23 group vs the ZT11 group. Since the comparison is across different time points, several genes are differentially expressed simply by virtue of the circadian rhythms or other time-dependent effects. To account for this, we normalized the differential expression in the IAV infected group at each time point for the control group from the same time point.

A p-value based two-way ANOVA analysis to extract interaction effects is generally considered unreliable with just three replicates per condition. However, achieving a meaningful ranking of genes that informs a powerful pathway enrichment analysis is sufficient for our purposes. A detailed study outlining our systematic approach is described to find the optimal value of pseudo-count (≈20) for an adjusted fold-change measure to rank genes by expression values in RNA-Seq data is currently under review as a methods based paper. The top 900 genes with difference of adjusted log10 fold-change greater than 0.67 (corresponding to about 5-fold change in the adjusted fold-changes) were used for pathway enrichment analysis. Pathway analyses were performed with Ingenuity IPA.

## Supplemental figures

**Table S1.**

| Score | Description |
| --- | --- |
| 1 | Percolated fur but no detectable behavior differences from untreated control mice |
| 2 | Mice with percolated fur and a huddle reflex but respond to stimuli (such as a tap on their cage) appropriately and are just as active upon handling as untreated control mice |
| 3 | Exhibit a slower response to a tap on the cage and that were passive or docile when handled but still curious when alone in a new setting |
| 4 | Exhibit lack of curiosity and little response to stimuli and that appear quite immobile. |
| 5 | Exhibit labored breathing and are unable or slow to right themselves after being rolled onto their backs (moribund) |
| 6 | Dead mouse. |

**Fig. S2:**
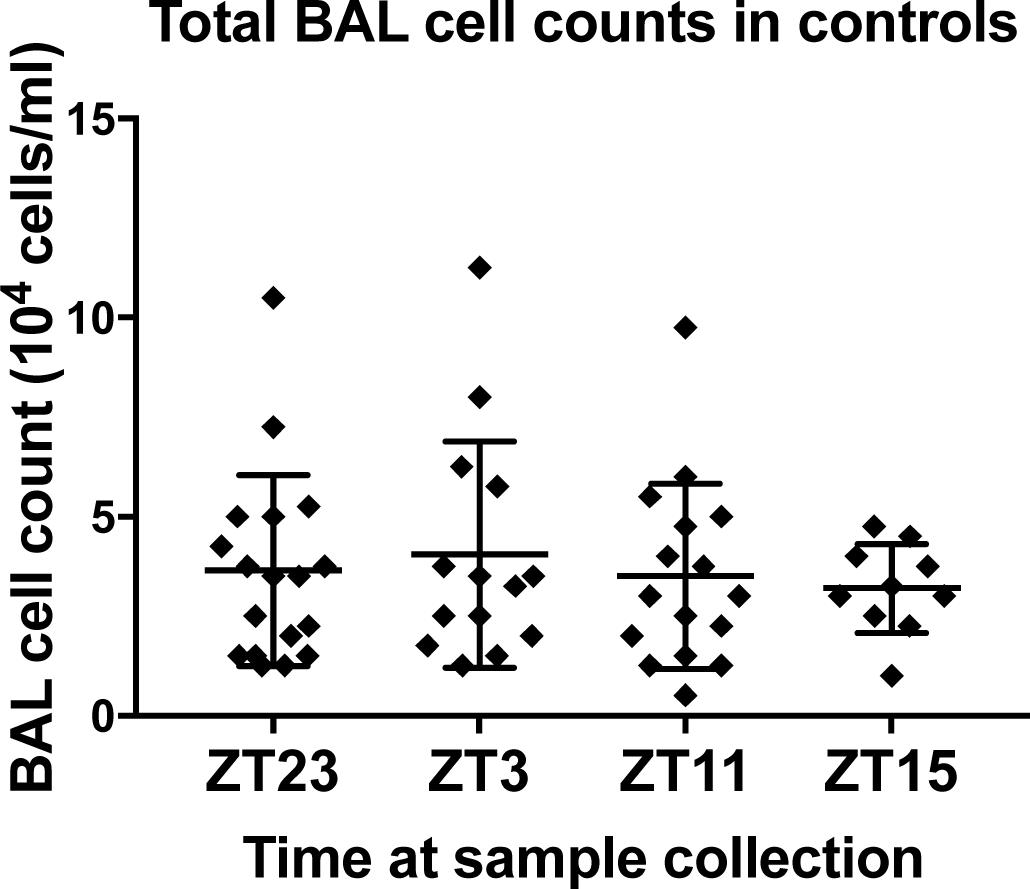

**Fig. S3:**
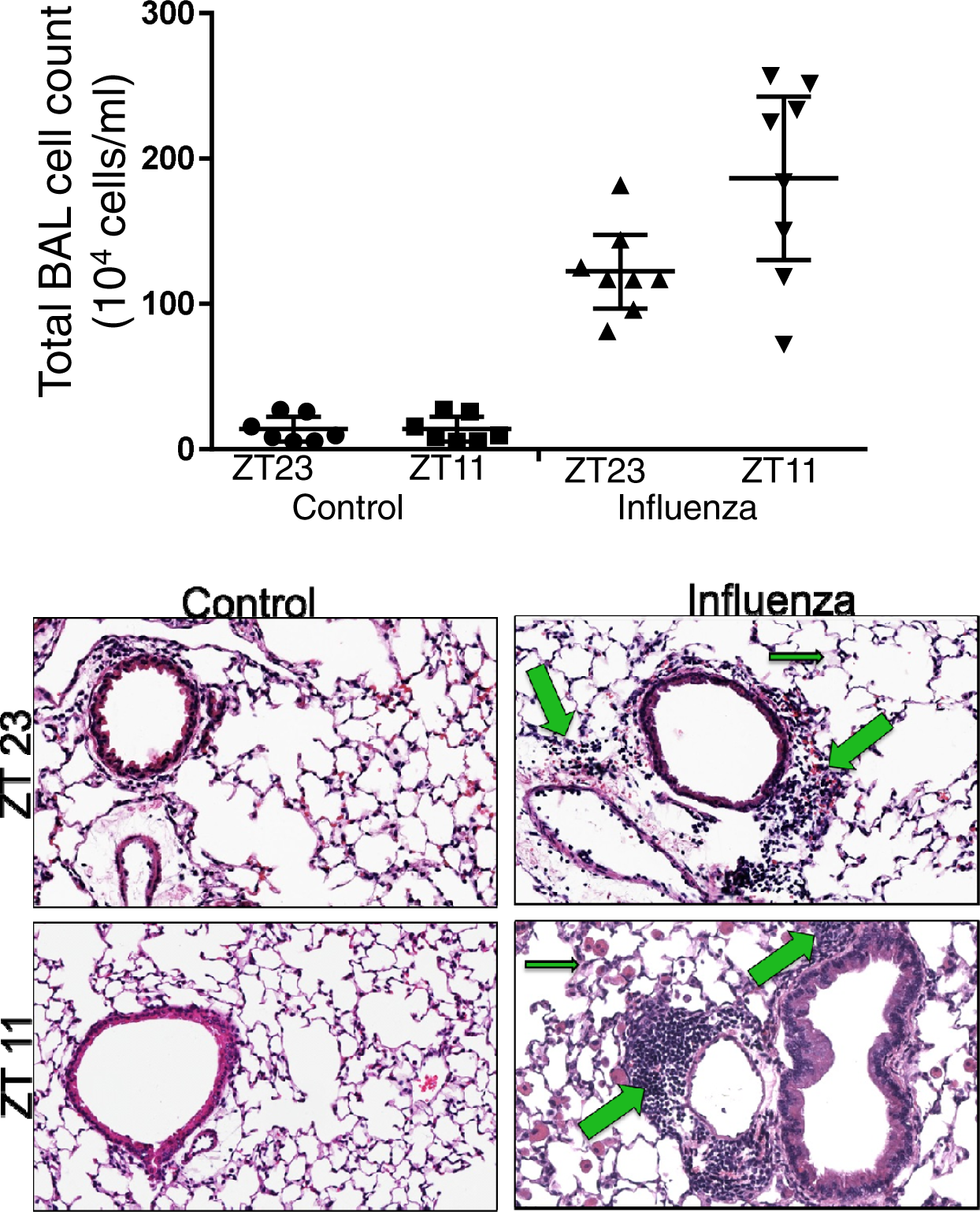

**Fig. S4:**
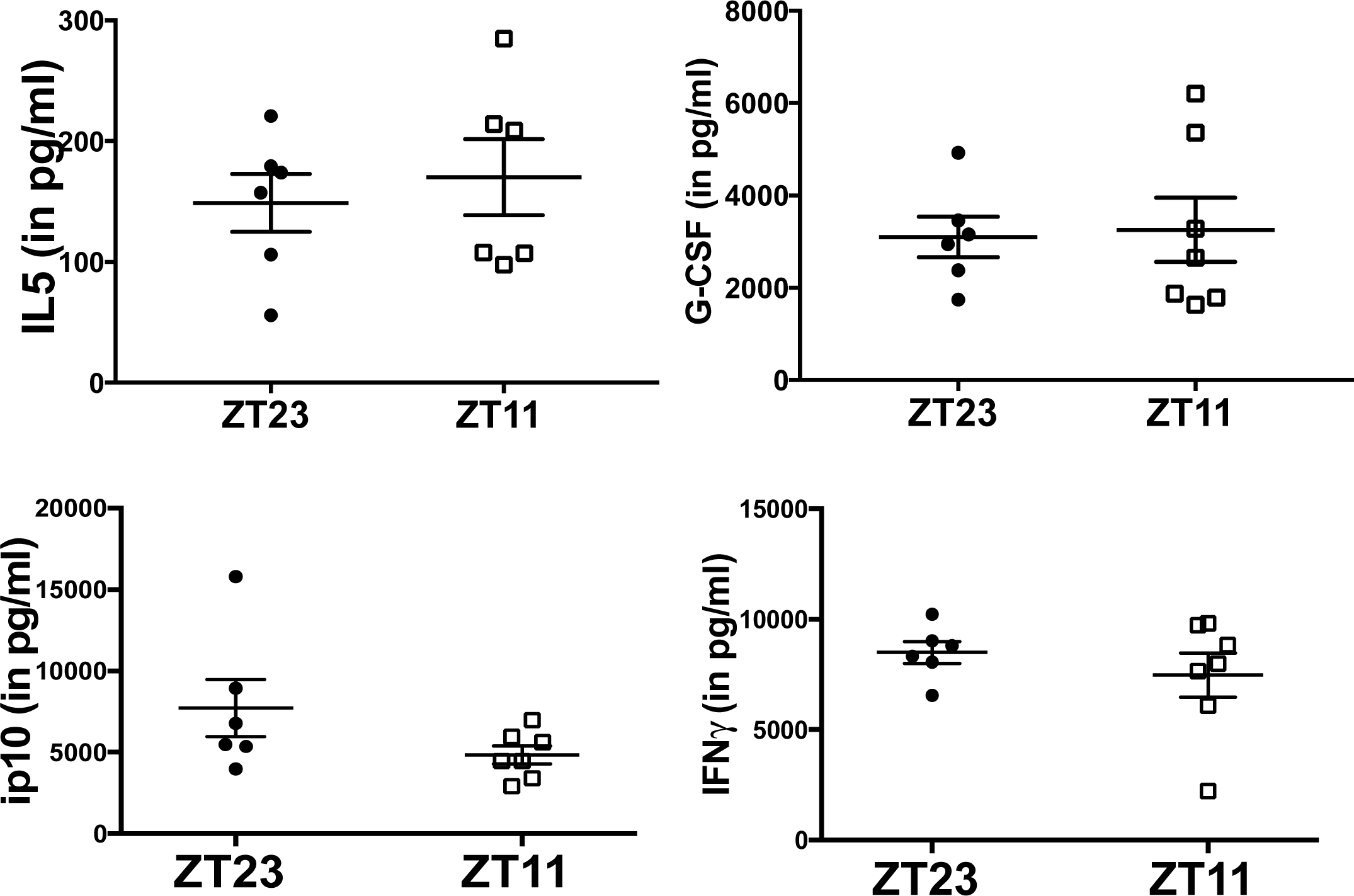

**Fig. S5:**
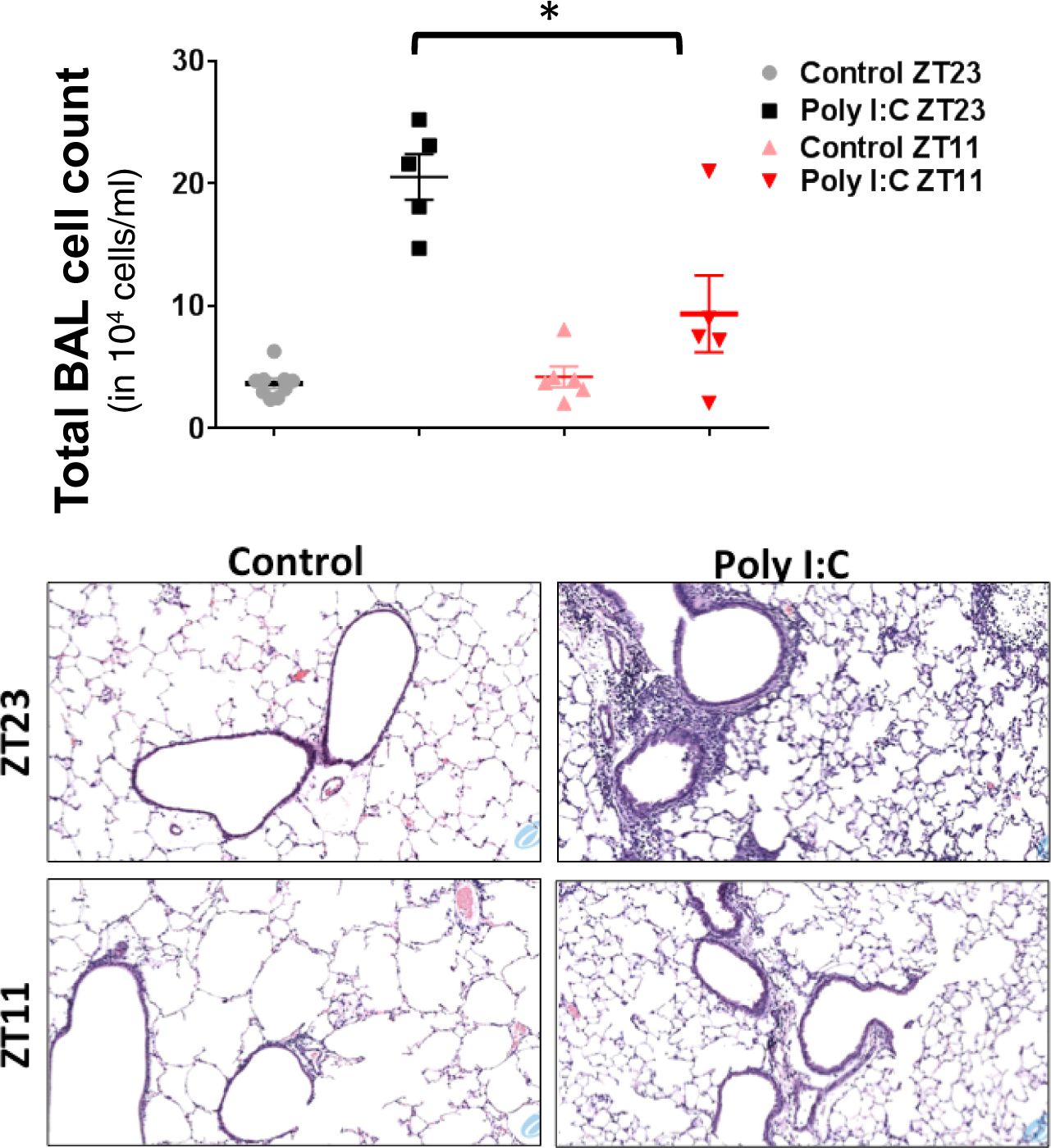
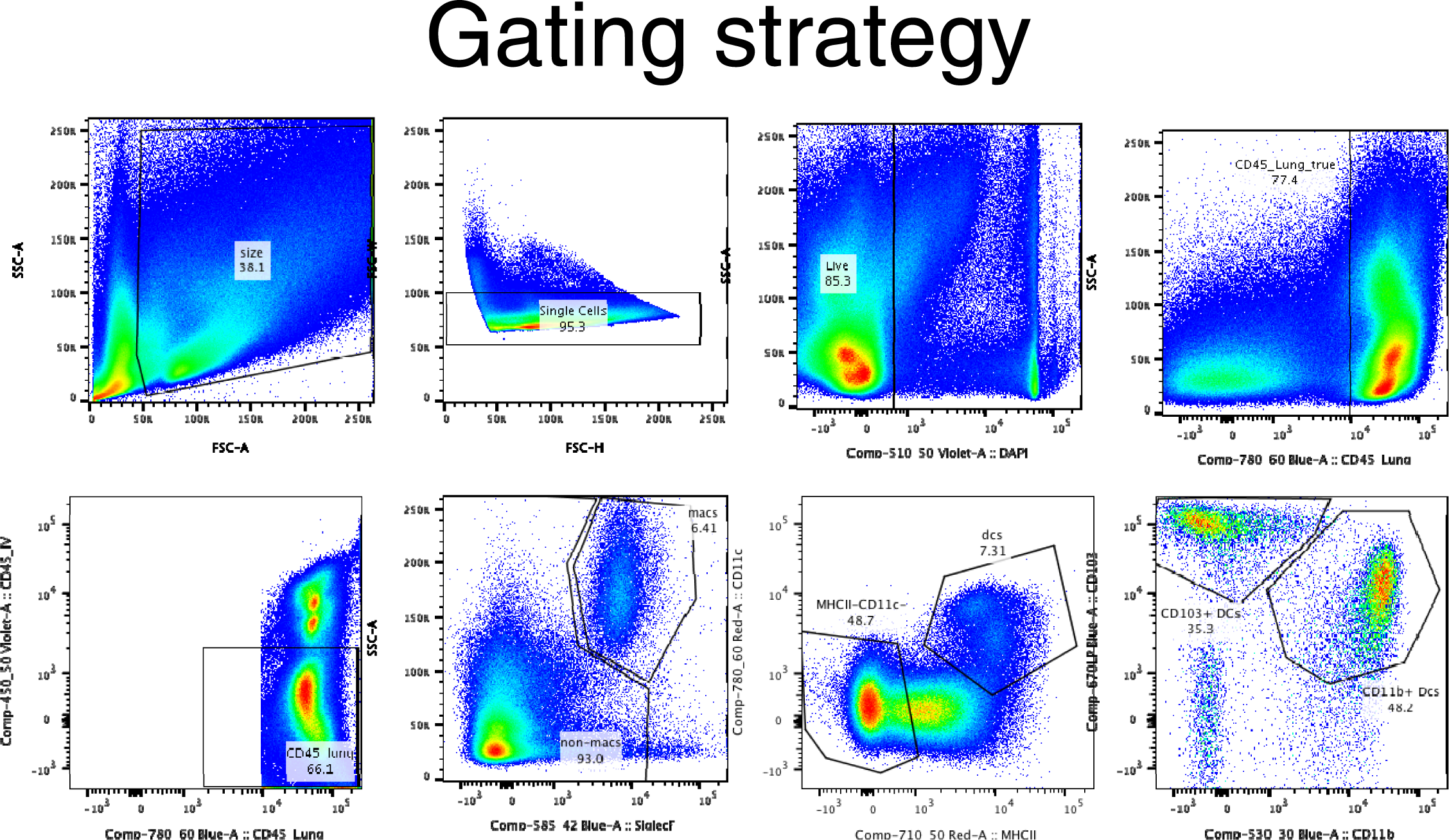

**Fig. S6:**
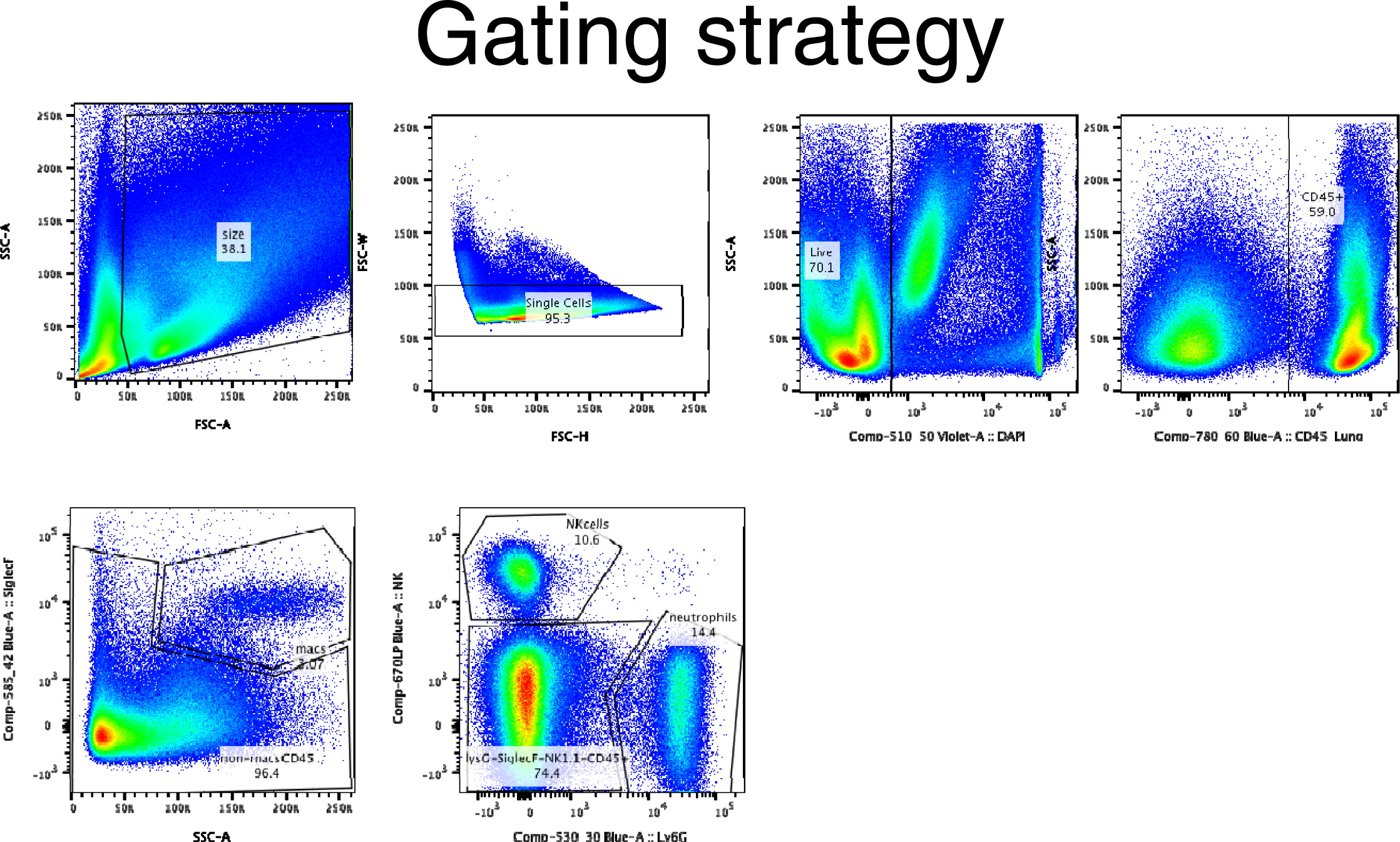

**Fig. S7:**
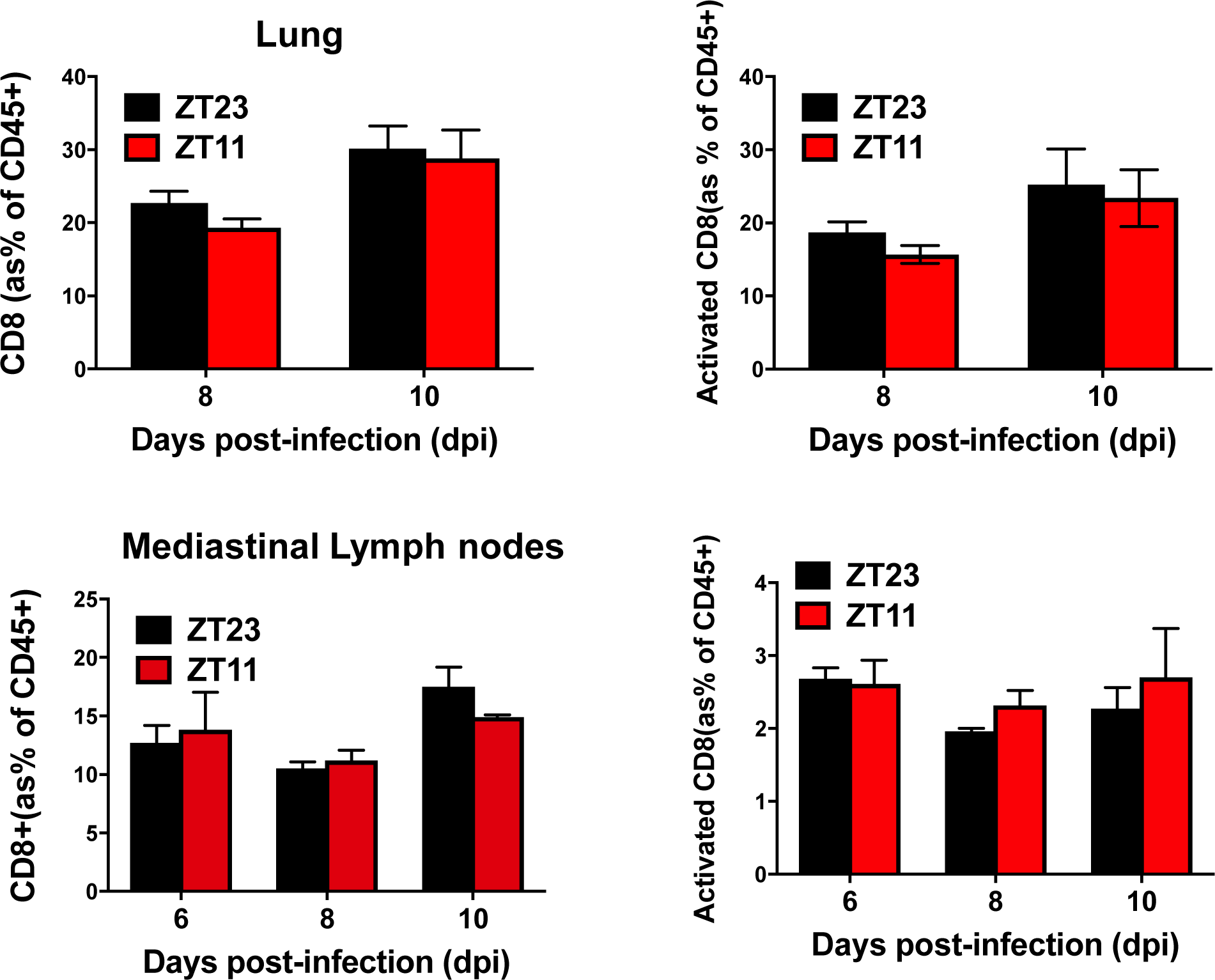

**Fig. S8:**
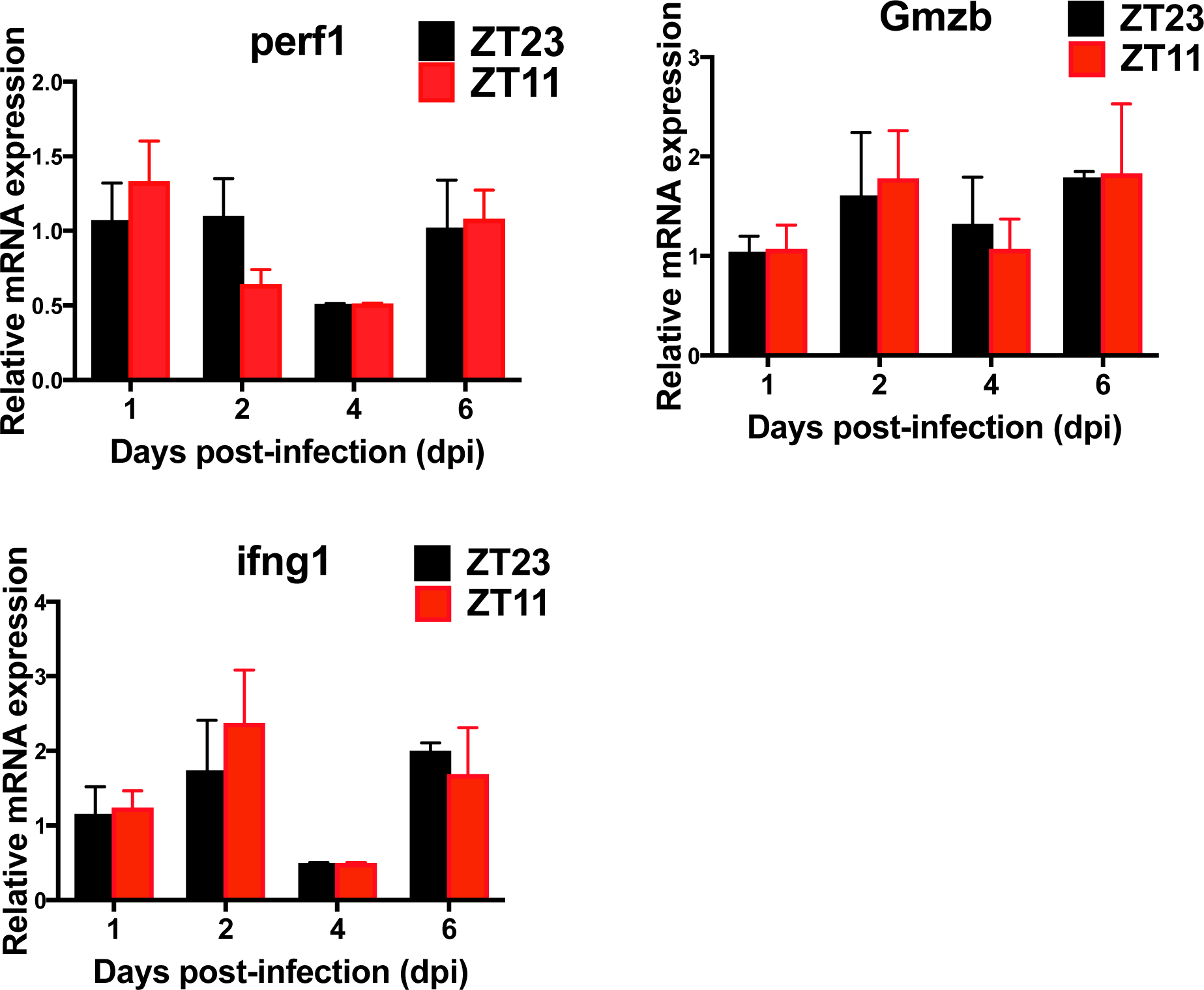
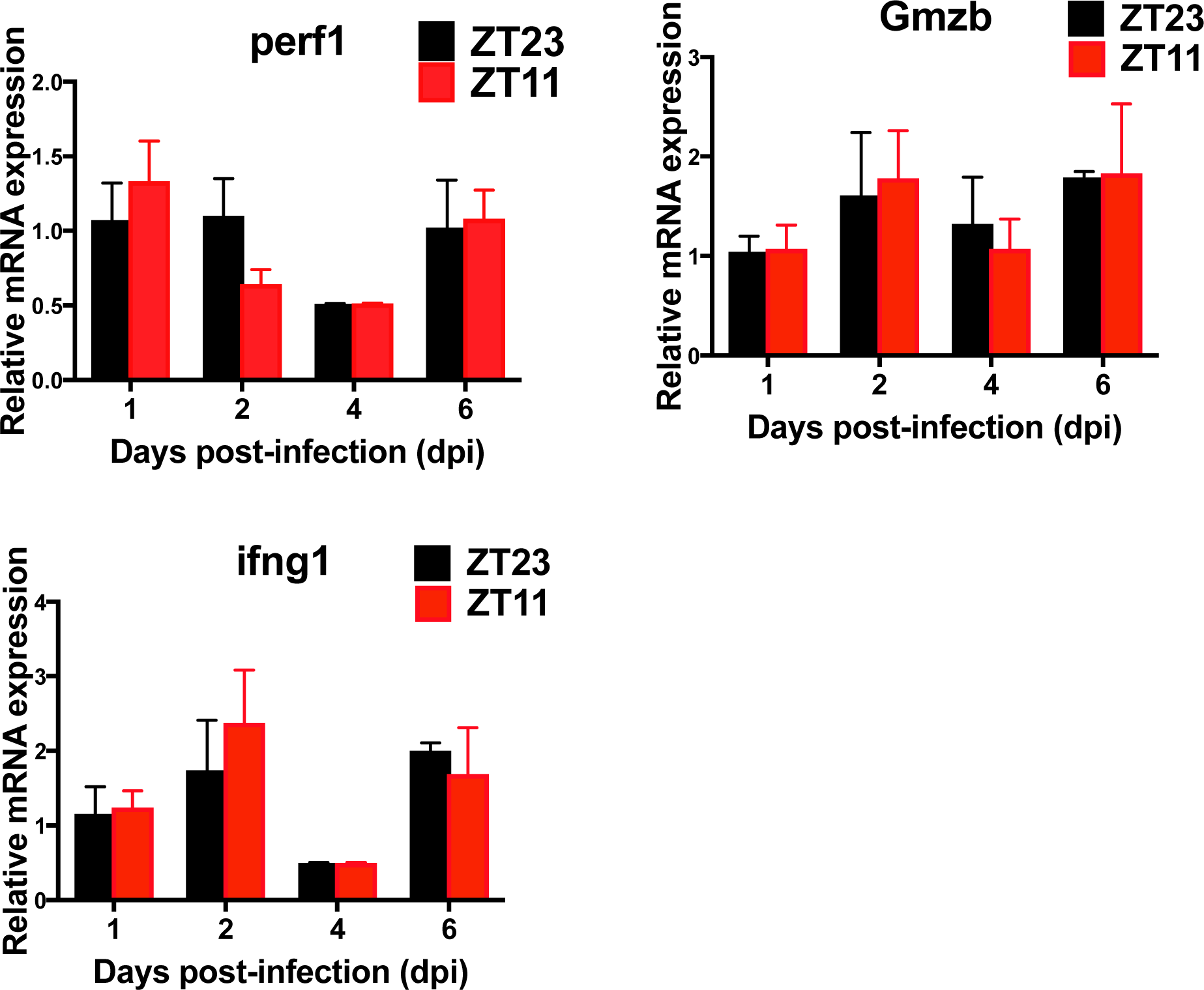

